# Integrative analysis identifies key molecular signatures underlying neurodevelopmental deficits in fragile X syndrome

**DOI:** 10.1101/606038

**Authors:** Kagistia Hana Utami, Niels H. Skotte, Ana R. Colaço, Nur Amirah Binte Mohammad Yusof, Bernice Sim, Xin Yi Yeo, Han-Gyu Bae, Marta Garcia-Miralles, Carola I. Radulescu, Qiyu Chen, Georgia Chaldaiopoulou, Herty Liany, Srikanth Nama, Prabha Sampath, Sangyong Jung, Matthias Mann, Mahmoud A. Pouladi

**Affiliations:** Translational Laboratory in Genetic Medicine, Agency for Science, Technology and Research (A*STAR), 8A Biomedical Grove, Immunos, Singapore 138648; Novo Nordisk Foundation Center for Protein Research, University of Copenhagen, 2200 Copenhagen N, Denmark; Singapore Bioimaging Consortium (SBIC), A*STAR, 11 Biopolis Way, Singapore 138667; Department of Physiology, National University of Singapore, Singapore 117597; Genome Institute of Singapore, Agency for Science, Technology and Research (A*STAR), 60 Biopolis Drive, 138672; Institute of Medical Biology, Agency for Science, Technology and Research (A*STAR), 8A Biomedical Grove, Immunos, Level 5, Singapore 138648; Department of Medicine, National University of Singapore, Singapore 117597

## Abstract

Fragile X syndrome (FXS) is an incurable neurodevelopmental disorder with no effective treatment. FXS is caused by epigenetic silencing of *FMR1* and loss of FMRP expression. To investigate the consequences of FMRP deficiency in the context of human physiology, we established isogenic *FMR1* knockout (*FMR1*KO) human embryonic stem cells (hESCs). Integrative analysis of the transcriptomic and proteomic profiles of hESC-derived FMRP-deficient neurons revealed several dysregulated pathways important for brain development including processes related to axon development, neurotransmission, and the cell cycle. We functionally validated alterations in a number of these pathways, showing abnormal neural rosette formation and increased neural progenitor cell proliferation in *FMR1*KO cells. We further demonstrated neurite outgrowth and branching deficits along with impaired electrophysiological network activity in FMRP-deficient neurons. Using isogenic *FMR1*KO hESC-derived neurons, we reveal key molecular signatures and neurodevelopmental abnormalities arising from loss of FMRP. We anticipate that the *FMR1*KO hESCs and the neuronal transcriptome and proteome datasets will provide a platform to delineate the pathophysiology of FXS in human neural cells.

## Introduction

Fragile X syndrome (FXS) is the most common genetic cause of intellectual disability and autism spectrum disorder (ASD) [1]. In addition to cognitive impairment, individuals with FXS often exhibit seizures, hypersensitivity, impulsivity, and social anxiety [2]. FXS is caused by an expansion of CGG trinucleotide repeats in the 5’ UTR of the *FMR1* gene, located on chromosome X, which results in specific hypermethylation of *FMR1* and silencing of FMRP, its encoded protein [3]. FMRP, a brain-enriched RNA binding protein, has been shown to regulate the translation of as many as 800 mRNAs in neurons by stalling ribosomes [4]. Loss of translational regulation of these mRNAs, whose products are involved in a wide range of neurodevelopmental and neuronal processes, is thought to underlie the pleiotropic molecular and clinical manifestation of FXS [5]. Despite continued therapeutic efforts and several clinical trials, no effective treatment has been developed to date [6].

The identification of individuals exhibiting a spectrum of FXS clinical phenotypes who carry mutations in *FMR1* that disrupt FMRP’s RNA binding activity have provided strong support for loss of function as the underlying cause of FXS [7–9]. This discovery further validated the use of *FMR1* knockout (KO) animal models to investigate the pathogenesis of FXS [10]. *FMR1* KO mice exhibit a number of phenotypes reminiscent of symptoms seen in individuals with FXS such as enlarged testes (macroorchidism), increased susceptibility to seizures and sensory hypersensitivity, hyperactivity, as well as perseveration and repetitive behaviours [10]. A number of molecular and synaptic defects have also been identified in rodent models of FXS [11], including abnormalities in dendritic spine morphology [12], protein synthesis [13], and neurotransmission [14] which, combined with the neurological deficits, have paved the way for the discovery and interrogation of novel targets for therapeutic intervention [6].

Human pluripotent stem cells (hPSCs) have emerged in recent years as a powerful tool to overcome the inaccessibility of the brain and to explore the underlying mechanisms of neurological diseases [15]. Studies using hPSCs to model neurodevelopmental disorders such as Rett [16], Down [17], Angelman [18], and Timothy syndrome [19] have begun to elucidate the neurodevelopmental abnormalities and pathogenic mechanisms associated with these disorders in the context of human physiology. For FXS, studies using hPSC-derived neurons have begun to identify disease-associated defects including abnormal morphologies as well as aberrant synaptic function [20–26]. However, a caveat of studies published to-date is the use of FXS and control hPSC lines with different genetic backgrounds. Here, we describe the generation of isogenic *FMR1* knockout (*FMR1*KO) human embryonic stem cell (hESC) lines and their use to investigate the pathophysiology of FXS in the context of human neural cells.

## Results

### Generation of isogenic *FMR1* knockout (*FMR1*KO) hESCs using CRISPR/Cas9

Isogenic pluripotent stem cells are important tools to model genetic disorders in the context of a common genetic background while working in cell types of interest. To generate *FMR1*KO hESCs, we utilized CRISPR/Cas9 nucleases targeting exon 3 of the *FMR1* gene in the male H1 hESC line (**Figure S1a**). We first evaluated the on-target activity of the *FMR1*-targeting sgRNAs in HEK293 cells using the Surveyor assay and confirmed cleavage for both sgRNAs (**Figure S1b**). Following electroporation of plasmids into the control hESCs, clones were screened for indels in *FMR1* using the Surveyor assay (**Figure S1c**). Eight clones with indels in *FMR1* out of approximately 48 colonies (17% efficiency) were obtained, of which two clones with 8 and 17 base pair (bp) deletions were selected for further characterization, based on their predicted amino acid truncation (**Figure 1a, Figure S1d**). PCR amplification for Cas9 in both *FMR1*KO lines showed absence of product at the expected size (2,045 bp), indicating no integration of the Cas9 transgene in the targeted clones (**Figure S1e**). We termed these two clones *FMR1*KO1 (8 bp deletion) and *FMR1*KO2 (17 bp deletion).

**Figure 1.**
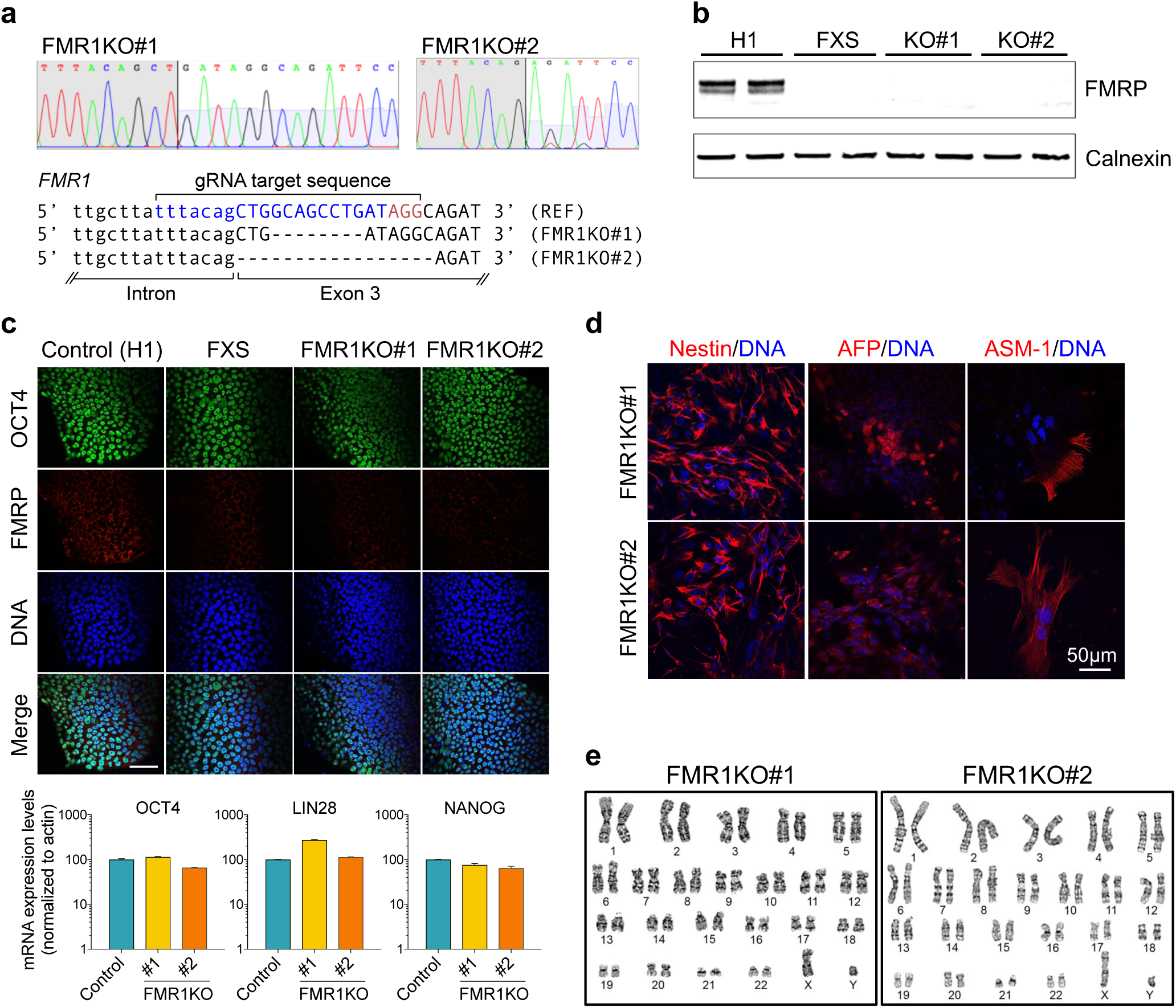
Generation of *FMR1*-/y H1 (*FMR1*KO) hESCs using CRISPR-Cas9 system. (**a**) Confirmation of indel and purity by Sanger sequencing. Schematic diagram showing the position of sgRNA sequence and indels generated in *FMR1*KO#1 and *FMR1*KO#2. (**b**) Immunoblot analysis showing absence of FMRP expression in FXS, *FMR1*KO1, and *FMR1*KO2. FMRP (80 kDa) and Calnexin (67 kDa). (**c**) *FMR1*KO hESCs express the indicated pluripotency markers as shown by immunostaining and qRT-PCR. Values shown as mean ± SD, based on n = 3 replicates per genotype. (**d**) *FMR1*KO hESCs can give rise to three germ layers: ectoderm (Nestin), mesoderm (ASM-1), and endoderm (AFP). (**e**) *FMR1*KO hESCs maintained a normal karyotype demonstrated by G-banding analysis.

To characterize the *FMR1*KO hESC lines and further interrogate neural phenotypes underlying FXS, we included a previously described FXS hESC line [27, 28] in our assessments. Consistent with FXS hESCs, we observed no expression of FMRP in *FMR1*KO hESCs by immunoblotting using two different antibodies that bind to the N-terminal region and the KH domains of FMRP (**Figure 1b, Figure S1f**) and immunofluorescence imaging (**Figure 1c**). Furthermore, proteome MS analysis confirmed the absence of FMRP peptides in the FXS and *FMR1*KO lines (**Figure S1g**). Loss of FMRP did not appear to compromise the pluripotency of the isogenic *FMR1*KO hESCs as indicated by the uniform expression of pluripotency markers (OCT4, LIN28 and NANOG) (**Figure 1c**). We next examined the ability of isogenic hESC lines to differentiate into cells of the three germ layers via a standard embryoid body formation protocol. The ability to differentiate in vitro was confirmed by the presence of ectoderm (Nestin), mesoderm (atrial siphon muscle-1, ASM-1), and endoderm (alpha feto-protein, AFP) markers (**Figure 1d**). Importantly, no chromosomal aberrations were introduced by the gene targeting process and the *FMR1*KO hESCs maintained a normal karyotype (**Figure 1e**).

### Transcriptome and proteome analyses reveal cellular pathways altered in FMRP-deficient neurons

To further investigate neural deficits caused by loss of FMRP, we differentiated control, *FMR1*KO, and FXS hESCs into neurons using a previously described protocol [29] (**Figure S2a**). By day 37, flow cytometry analysis of the differentiated cultures, using an established panel of cell surface markers of neural and glial cells [30], showed that the proportion of neural versus glial cells derived from the different lines was comparable (**Figure S2b**). Further characterization of the resulting neuronal population showed them to be comprised mostly of MAP2/TUJ1-positive glutamatergic neurons (∼65-80%) with a lower proportion of GABAergic neurons (∼20%) (**Figure S2c,d**).

To gain insights into the cellular processes disrupted by loss of FMRP, we performed RNA sequencing on day 37 neurons differentiated from control (H1) and isogenic *FMR1*KO hESCs, as well as FXS hESCs (n = 4 for each cell line). Principal component analysis (PCA) of the transcriptional profiles showed tight clustering of biological replicates per genotype (**Figure 2a**). We identified a substantially higher number of differentially expressed genes (DEGs) in *FMR1*KO neurons versus isogenic control neurons compared with FXS versus control, as depicted in the volcano plots (**Figure 2b,c**). In total, we identified an overlap of 3,110 DEGs from FXS and *FMR1*KO neurons that were altered compared with controls, of which 1,525 genes were downregulated and 1,585 were upregulated (**Figure 2d-g**). Further statistical analysis with different threshold of log2 fold change reduced the number of DEGs substantially, indicating that most of the DEGs (83-97%) had smaller fold changes (|log2FC|> 0). Separation of the differentially expressed genes by log2 fold change (FC) showed that 17%, 7%, 3% of the DEGs have a |log2FC|> 1, > 1.5, and > 2, respectively (**Figure 2f**). Clustering of the 3,110 DEGs showed a high correlation between expression levels in FXS and *FMR1*KO samples supporting the current selection of genes of interest (**Figure 2g**). Functional annotation of DEGs shared between *FMR1*KO and FXS using gene ontology (GO) analysis and network visualization showed enrichment of a number of GO terms, including those related to neuron differentiation, and neurodevelopment, neurogenesis, neurotransmission for downregulated genes, and RNA processing and transport, translation, and cell cycle processes for upregulated genes (**Figure 2h, Figure S3a**). Among the neuronal differentiation GO categories, we identified genes involved in axon guidance, neurite outgrowth, and cell adhesion, such as *DSCAM, GAP43*, and *PTPRT*. To validate the changes between neuronal lines detected by RNA-seq, we confirmed these alterations by qRT-PCR (**Figure S3b**).

**Figure 2.**
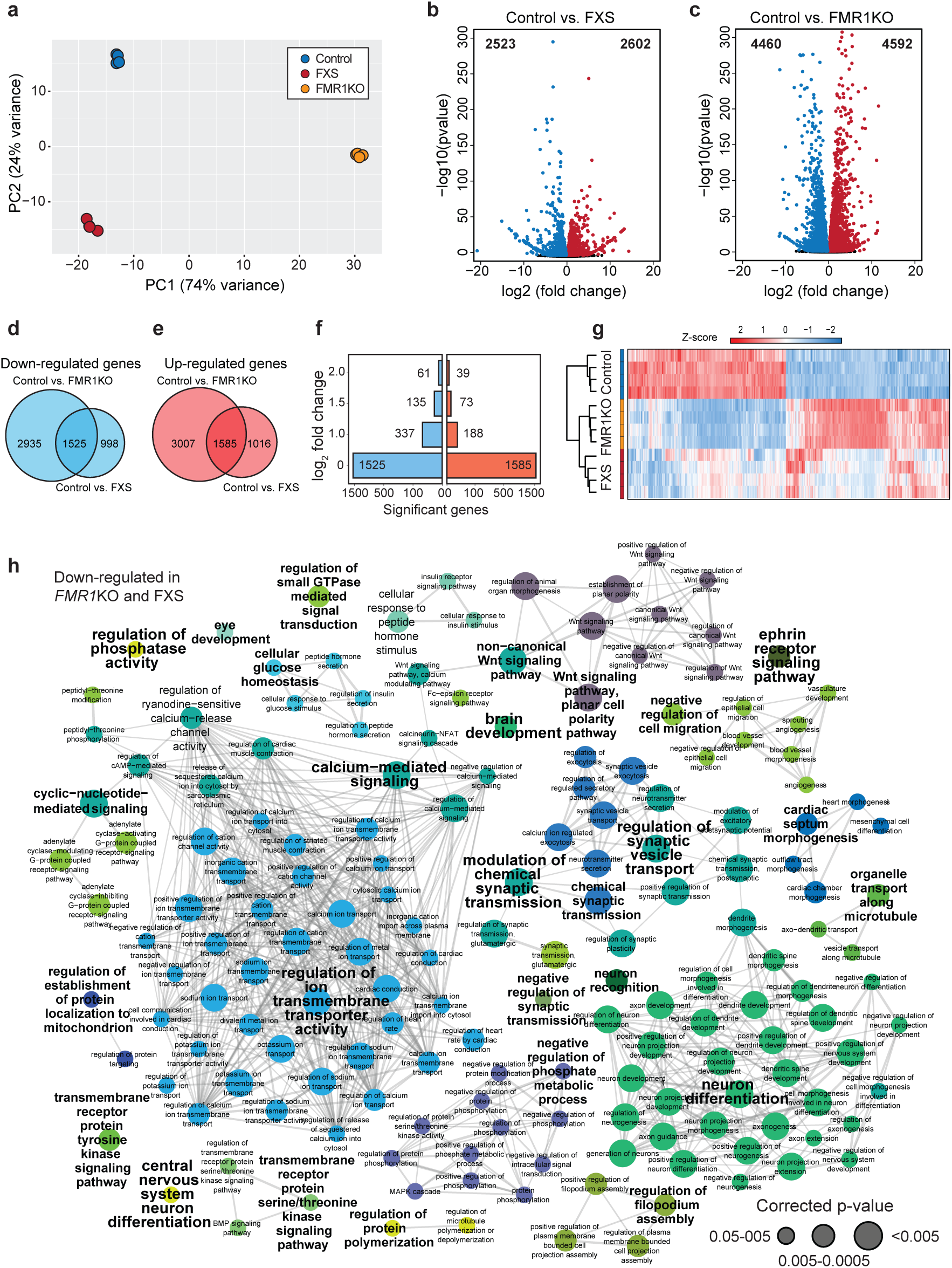
Global transcriptional changes in FMRP-deficient neurons. (**a**) Principal component analysis (PCA) shows tight clustering of replicates for each genotype; (**b,c**) Volcano plots of -Log10 (p-value) versus the Log2 (fold change) of transcript levels for all genes. Relative to control, significantly downregulated genes are shown in blue and upregulated genes are shown in red; (**d,e**) Venn analysis showing genes that are similarly downregulated (**d**) and upregulated (**e**) between control vs FXS and control versus *FMR1*KO; (**f**) bar plot showing the number of significant changes dependent on the cut-off in log2 fold changes, (**g**) Heatmap of differentially expressed genes that are common to FXS and *FMR1*KO neurons; (**h**) Functional annotation of common downregulated genes in FXS and *FMR1*KO neurons; functional annotation of common upregulated genes can be found in **Figure S3a**.

We compared our list of commonly identified genes in FXS and *FMR1*KO with two previously published transcriptome datasets of FXS neurons [23, 31], where we found a solid overlap of the identified genes (**Figure S4a**). From the shared genes between the three studies, we found that approximately 50% of our significantly regulated genes were also regulated in either the Boland et al. or the Halevy et al. study. (**Figure S4b** and **Table S2**). Importantly, of our regulated DEGs, we found significant association with 239 Simons Foundation Autism Research Initiative (SFARI) genes, a database of genes linked to autism [32] (**Figure S4c,d** and **Table S2**) (|log2FC |>0, p=6.35 × 10^-10^ and |log2FC|>1, p=7.82 × 10^-3^). We also observed significant association for DEGs with |log2FC |> 0 for FMRP substrates (246 genes, p=1.1 × 10^-29^) (**Figure S4c** and **Table S2**). Together, our findings support that loss of FMRP results in transcriptional dysregulation, which alters processes involved in central nervous system development, such as neuronal differentiation, neurogenesis, and cell cycle regulation.

To map the FXS proteome, we collected neurons on day 37 of differentiation, the same time-point used for the transcriptome analysis, and subsequently profiled protein expression changes by MS analysis. To establish a baseline for the technical and biological quality, we first compared protein and peptide numbers and evaluated the reproducibility. From this, we identified a total of 5007 proteins, where 4210 were selected for further investigations after stringent filtering, and compared protein and peptide identification across the 3 genotypes (**Figure S5a,b**). The majority of the selected proteins in our study were shared between all three genotypes (**Figure S5c**). Furthermore, we observed strong reproducibility (Pearson correlations ranging from 0.97-0.98) between biological replicates (**Figure S5d**). The high reproducibility in our analysis was further supported by a PCA demonstrating a strong separation of the three genotypes (**Figure 3a**), supporting that our proteomic analysis provides a sound foundation for genotypic comparison. Next, we performed a system-wide comparison in more detail comparing FXS and *FMR1*KO to control, respectively. (**Figure 3b,c** and **Figure S5e,f**). Similarly to the results from the RNA-seq statistical analysis, the vast majority of significantly regulated proteins depicts smaller log2 fold changes (|log2FC |> 0) with only 17% and 13% of down- and upregulated proteins, respectively, showing log2 fold changes beyond 1 (|log2FC|> 1) (**Figure S5e**). A hierarchically-clustered heatmap shows grouping of the samples according to the expected genotypes, whereas column values suggest a closer correlation between FXS and *FMR1*KO expression levels than with expression levels in Control samples (**Figure S5f**). We identified several protein changes with a total of 577 and 2198 proteins exhibiting increased or decreased expression in FXS and *FMR1*KO compared to control, respectively, as depicted in volcano plot (**Figure 3b,c**).

**Figure 3.**
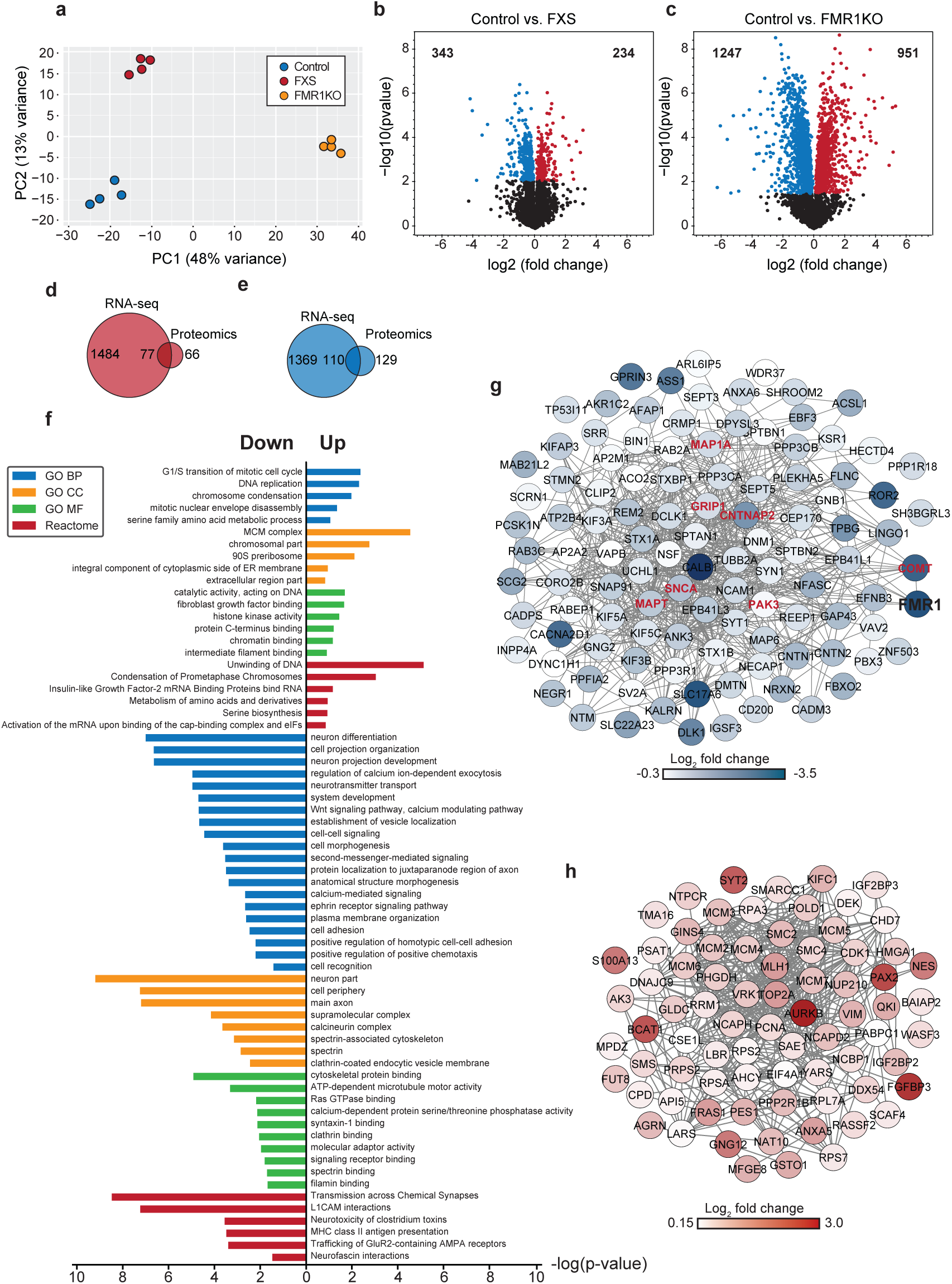
Global proteomic changes in FMRP-deficient neurons. (**a**) Principal component analysis (PCA) shows clustering of replicates for each genotype; (**b,c**) Volcano plots of -Log10 (p-value) versus the Log2 (fold change) of protein levels for all proteins. Relative to control, significantly downregulated proteins are shown in blue and upregulated proteins are shown in red; (**d,e**) Venn analysis showing common significant changes shared between FMR1KO and FXS as well as between the RNA-seq and proteomics data. 77 upregulated proteins/genes (**d**) and 110 downregulated (**e**) proteins/genes; (**f**) Functional annotations for the significant common changes, where a subset (based on protein with lowest p-value per cluster group for GO terms BP, CC, and MF as well as Reactome) is displayed. (**g,h**) STRING protein-protein interactions network for the common significant upregulated (**g**) and downregulated (**h**) changes at both the RNA and protein level are illustrated. Proteins associated with *FMR1* are highlighted in red.

Next, we performed pathway enrichment analysis for significantly regulated proteins in FXS (343 downregulated and 234 upregulated proteins) and *FMR1*KO (1247 down-regulated and 951 upregulated proteins) samples, separately. In the same way as the RNA-seq data, the enriched GO terms for differentially expressed proteins in *FMR1*KO showed down-regulation of proteins involved in neuron development, neuron differentiation, neurotransmitter secretion, and regulation of calcium ion transport, whereas the upregulated proteins gave rise to GO terms including RNA processing and splicing and ribosome biogenesis. In addition, the GO terms for differentially expressed proteins in FXS showed enrichment in vesicle transport and synaptic signalling for downregulated proteins, and cell cycle processes, DNA replication and DNA metabolic processes for upregulated proteins. Among the downregulated proteins we observed several candidates such as *CNTNAP2* [33, 34], *GPRIN3, KIF5C* [35] and *CNTN1* previously associated with FXS and other neurodevelopmental disorders as well as neurodegenerative diseases.

To identify and summarize the common changes between in FXS and *FMR1*KO, we performed pathway enrichment analysis for differentially expressed proteins shared between FXS and *FMR1*KO (244 downregulated and 141 upregulated proteins). We condensed the data and visualized the GO terms for biological processes (GOBP) terms as two networks representing up- and downregulated common protein changes (**Figure S6a,b**). Interestingly for the networks, we found cellular processes based on downregulated proteins related to neurotransmitter secretion, synaptic transmission, transport, and dendrite development and DNA replication, gene expression, and cell cycle processes for upregulated proteins. Importantly, the data reveal many GO categories similar to those identified in the transcriptional enrichment analysis, highlighting the complementary of the proteomic and transcriptional analyses and providing further evidence linking loss of FMRP to functional perturbations in these pathways. This notion is further supported by the enrichment of both SFARI and *FMR1*/FMRP related proteins (**Figure S7a,b**). In the proteomics data set, we found significant association with 40 SFARI database genes and 57 FMRP genes (|log2FC |>0; p=8.32 × 10^-3^ and p=2.12 × 10^-7^, respectively) (**Figure S7a,b** and **Table S3**), which demonstrated that our findings at the RNA level translated to the protein level providing another layer of functional insight. Collectively, our findings highlight several biological processes and pathways altered in FMRP-deficient neurons of potential relevance to the pathogenesis of FXS.

Lastly, to combine the knowledge gathered from the RNA-seq and proteomics analyses, we explored the overlap of significant changes between the two data sets (**Figure S8a**). First, we compared the expression correlation between the two data sets and found a good correlation between the RNA and protein expression (R2=0.42; p=8.6 × 10^-171^, R2=0.44; p=9.9 × 10^-190^, and R2=0.42; p=2.9 × 10^-171^, for control, *FMR1*KO, and FXS, respectively, **Figure S8b**). Notably, we found significant 187 genes and proteins in common between the two data sets, where 77 and 110 were significantly up- and downregulated, respectively (**Figure 3d-e**). The enrichment analysis on the common changes between the two data types (RNA-seq and proteomics) revealed several altered interesting pathways related to neurotransmitter transport, synaptic signalling, neuron differentiation, and brain development for down regulated proteins/genes (**Figure 3f**) as well DNA replication, mitosis, and cell cycle for upregulated proteins/genes. These findings clearly encapsulate the commonalities between the two data sets and demonstrate that the resources described in this manuscript can be used independently or in combination. Lastly, in order to better understand how these genes and proteins might interact with each other, we looked into protein-protein interaction networks using STRING database (string-db.org) (**Figure 3g,h**). The networks were plotted using the edge betweenness clustering algorithm to identify highly connected nodes. This method clusters together proteins, which are known to cooperate and have correlating gene function annotations [36]. From this analysis, we found six genes (*COMT, MAP1A, GRIP1, PAK3, SNCA, MAPT*, and *CNTNAP2*) to be closely associated with *FMR1*/FMRP. Interestingly, *SNCA* (alpha-synuclein), *MAPT* (Tau), *CNTNAP2* (Contactin associated protein-like 2), *PAK3* (P21 protein), and *COMT* (catechol-O-methyltransferase) have associations with Parkinson’s disease, Alzheimer’s disease, autism, intellectual disability, and schizophrenia, respectively. These findings were further supported by a search for curated gene-disease associations in DisGeNET database (www.disgenet.org). From our 187 common candidates, 53 downregulated and 28 upregulated genes were found to be associated with “Nervous System Diseases” and “Mental Disorders”.

### Loss of FMRP leads to abnormal neural rosette formation and increased neural progenitor proliferation

The transcriptional and proteomic analysis highlighted changes in neuronal development and neurogenesis; therefore, we sought to examine neural rosette formation following neural induction as a measure of early neurodevelopment in *FMR1*KO and FXS lines. We differentiated hESCs into neural rosettes using a previously described floating embryoid body method [37]. Neural rosettes appeared markedly smaller in the FXS and *FMR1*KO lines compared with the control line (**Figure 4a**). To quantify the size differences, we measured the area stained with ZO-1, a luminal neural rosette marker, and found it to be significantly reduced in *FMR1*KO and FXS lines compared with control (**Figure 4b**). Functional annotation of the upregulated genes in FMRP-deficient neurons showed enrichment of transcripts involved in mitosis and cell cycle-related processes. This prompted us to investigate whether cell proliferation is affected in FMRP-deficient cells. Labeling with BrdU and Ki67, two markers of cell proliferation, showed increased FXS and *FMR1*KO neural progenitor proliferation compared with control cells (**Figure 4c,d**).

**Figure 4.**
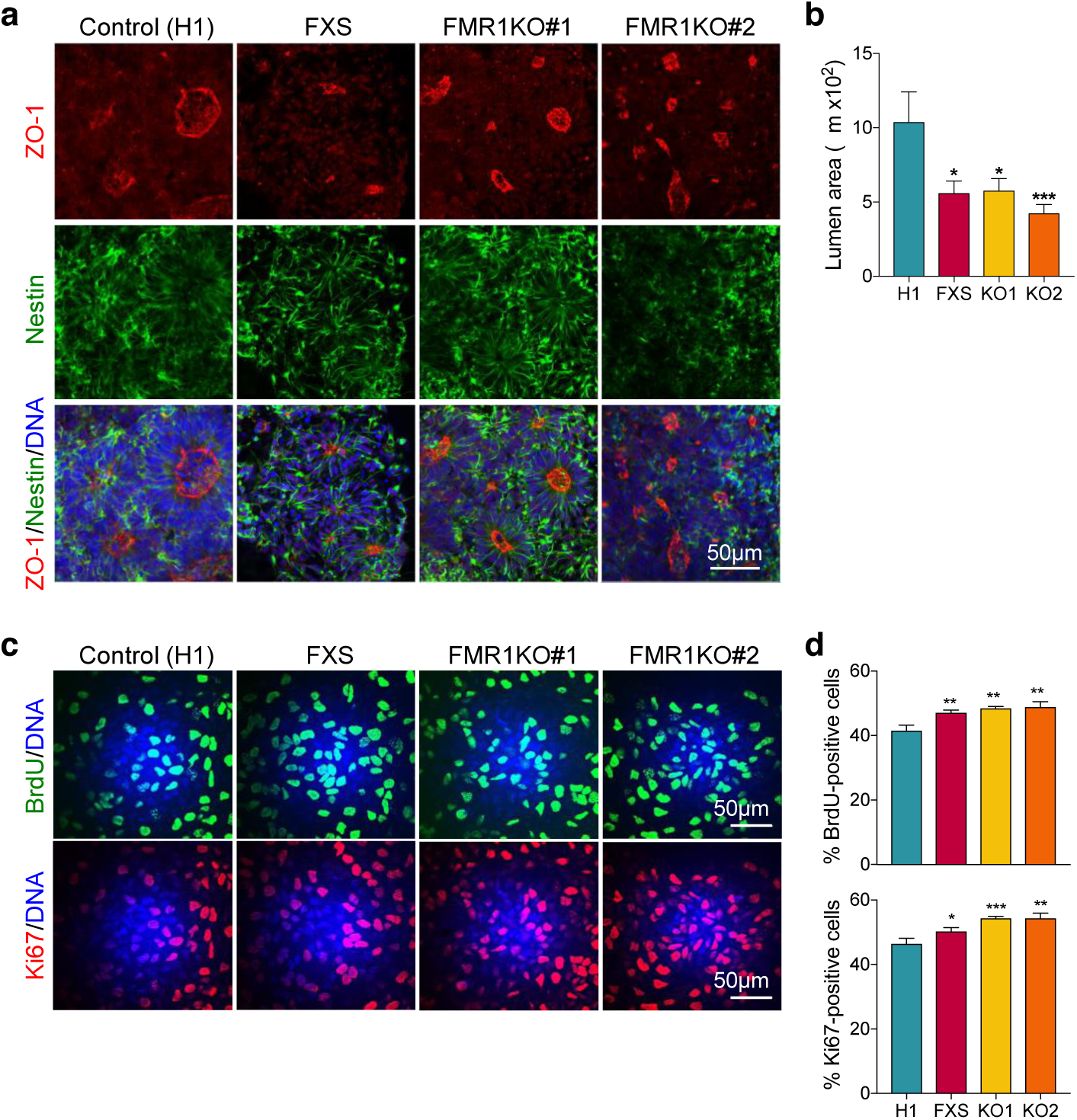
FMRP deficiency leads to abnormal neural rosette formation and increased neural progenitor proliferation. (**a**) Immunostaining shows neural rosette structures identified using Nestin and ZO-1 expression. (**b**) Quantification of lumen size area (based on ZO-1 positive staining). Lumen area was measured by ImageJ software. Values shown as mean ± SEM based on n = 4 biological replicates per genotype; *p < 0.05 and ***p < 0.001 compared to control (H1) was determined by one-way ANOVA with Tukey’s post-hoc test. (**c,d**) BrdU-labeling and Ki67 reveals increased proliferation in FXS and FMR1KO (KO1) neural progenitor cells compared with control (H1). Values shown as mean ± SEM based on blinded counting of 10 images from each of three biological replicates per genotype; *p < 0.05, **p < 0.01, and ***p < 0.001 compared with control (H1) was determined by one-way ANOVA with Fisher LSD post-hoc test.

### FMRP-deficient neurons exhibit neurite outgrowth deficits and abnormal network connectivity

Neurite outgrowth is an early neurodevelopmental process critical to the proper formation of axons and dendrites. Neurite outgrowth has been shown to be compromised in a number of intellectual disability and autism disorders, including FXS [22, 24, 38]. To evaluate this deficit in *FMR1*KO lines, we performed longitudinal tracking of neurite elongation in hESC-derived neurons using live-cell imaging. We found a striking reduction in neurite outgrowth and branching in *FMR1*KO and FXS neurons over time compared with control neurons (**Figure 5c,d**). These results demonstrate that isogenic *FMR1*KO neurons recapitulate this FXS-linked morphological deficit.

**Figure 5.**
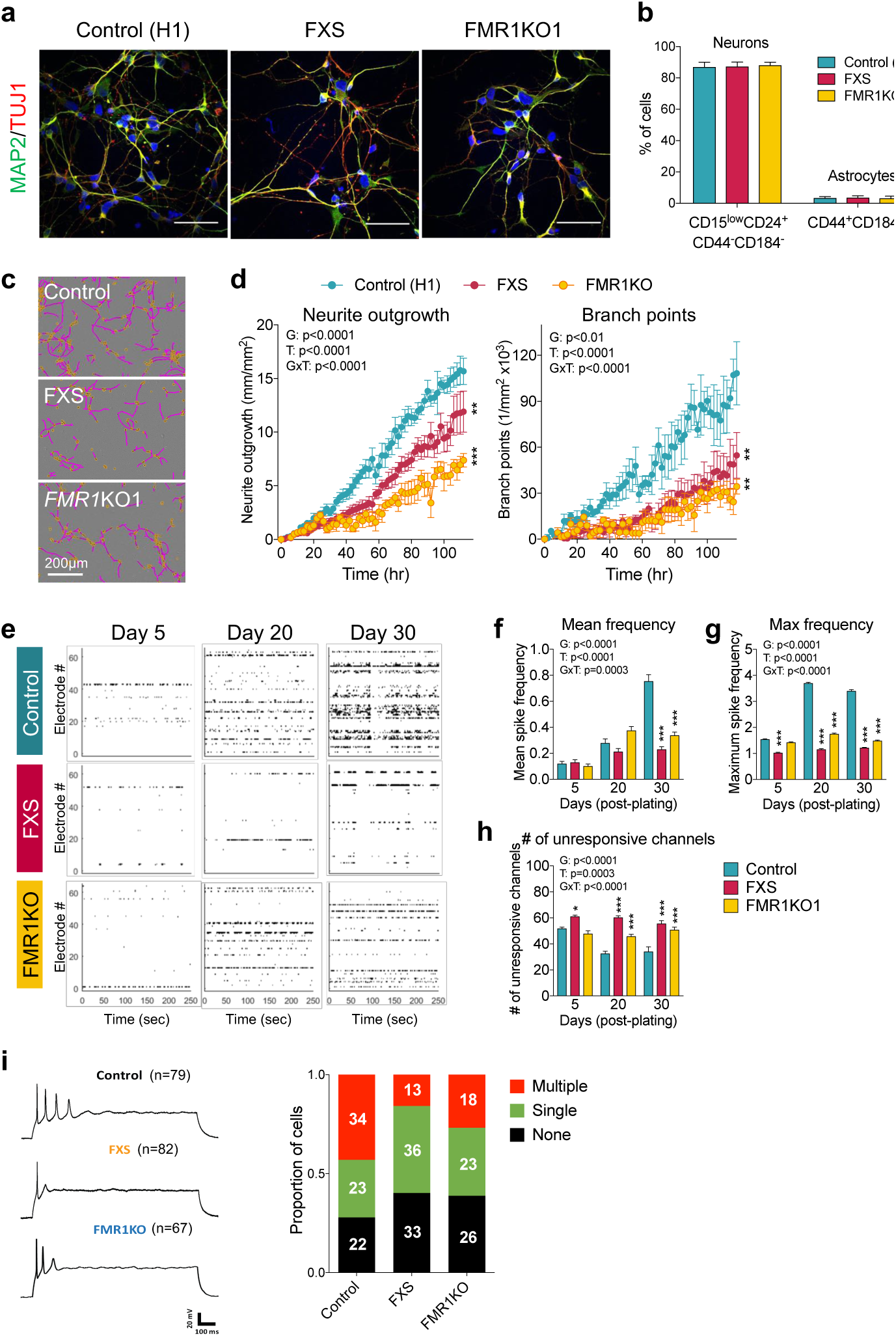
FMRP-deficient neurons exhibit shorter neurite length and aberrant network connectivity. (**a**) Immunostaining showing the expression of post-mitotic neuronal markers MAP2 and TUJ1 after 37–40 days of differentiation from hESC; scale-bar = 50 µm. (**b**) Flow cytometry analysis of cellular composition following hPSC neuronal differentiation using cell surface markers of neurons and glia. (**c**) Representative images of neurite phenotype segmentation. Neurites are labeled in yellow, and the neuronal cell bodies are labeled in pink/purple. (**d**) Neurite outgrowth and branching measurements. Values shown as mean ± SD from 6 biological replicates. (**e-f**) Analysis of neuronal network activity using multi-electrode array (MEA) recordings. n = 6 per genotype. (**e**) Raster plots of spike time stamps indicating neuronal spontaneous activity as measured by MEA recordings. (**f**) Mean firing (spike) frequency. (**g**) Maximum firing (spike) frequency. (**h**) Number of unresponsive channels; *p < 0.01, **p < 0.01, and ***p < 0.001 compared to control (H1) was determined by two-way ANOVA with Tukey’s post-hoc test. G, genotype: T, time. (**i**) Analysis of intrinsic electrophysiological neuronal properties by patch clamp.

### FMRP-deficient neurons show abnormal electrophysiological network connectivity

The reduced neurite outgrowth in FMRP-deficient neurons led us to investigate electrophysiological neuronal network connectivity. We evaluated extracellular spontaneous activity using longitudinal multi-electrode array (MEA) recordings, which enable detection of action potentials (spikes) based on changes in field potential [39]. MEA recordings were performed over a 30-day period starting at day 37 of neuronal differentiation. Representative raster plots depicting neuronal firings (spikes) over a 250-sec period of continuous recording on days 5, 20, and 30 are shown in **Figure 5e**. Spontaneous action potentials were detected for all groups, confirming successful derivation of functional neurons from all three genotypes. However, analysis of spike frequencies revealed profound deficits in the mean and maximum spike frequencies for *FMR1*KO and FXS neurons compared with control neurons (**Figure 5f-h**), suggesting altered neuronal network connectivity as a result of FMRP deficiency. As differences in MEA activity can also reflect intrinsic neuronal differences, we analyzed the basic properties of action potential (AP) firing in FMRP-deficient and control neurons using patch-clamp recordings (**Figure 5i** and **Table S4**). While there were no differences in the proportion of neurons with a single or no AP between FXS and *FMR1*KO (69 of 82 for FXS and 49 of 67 for *FMR1*KO, p > 0.05, Chi2 test), the proportion of such neurons was significantly larger when compared with control neurons (45 of 79 for control, p<0.001 compared with FXS, p<0.05 compared with *FMR1*KO, Chi2 test). These results suggest that differences in the intrinsic properties of FMRP-deficient neurons may also contribute to the deficits in MEA activity observed.

## Discussion

Isogenic models represent well-controlled experimental systems in which cellular and molecular abnormalities arising from genetic disorders can be readily attributed to the genetic lesion under study [40]. Here, we describe the generation and characterization of isogenic *FMR1*KO human embryonic stem cells as a model of FXS. We illustrate alterations of several biological pathways important to brain development and function in FMRP-deficient neurons at both the RNA and protein level. We functionally validate alterations in a number of these pathways, showing phenotypic deficits including abnormal neural rosette formation and increased neural progenitor cell proliferation. We further demonstrate neurite outgrowth and branching deficits in FMRP-deficient neurons along with impaired electrophysiological neuronal connectivity. Our findings reveal key molecular signatures and neurodevelopmental abnormalities arising from loss of FMRP.

We find that neurons lacking FMRP show dysregulation of genes and proteins related to diverse cellular and neuronal processes. It is interesting to note that there was a greater number of differentially expressed genes and proteins between the isogenic lines (*FMR1*KO versus control) compared with the non-isogenic ones (FXS versus control). The use of isogenic lines minimizes the potential for not only false positives but also false negatives, and it is possible that a large number of false negatives in the FXS versus control comparison due to differences in the genetic background resulted in the smaller number of differentially expressed genes and proteins in that group.

The cellular and neuronal processes dysregulated in FMRP-deficient neurons include nervous system development, neuronal differentiation, cell cycle progression, as well as neurotransmission. We were able to corroborate a number of these changes functionally. Firstly, we discovered that the formation of neural rosettes is abnormal in the absence of FMRP. Neural rosettes are self-assembling structures considered surrogates of early neurulation and neural tube formation [41, 42]. Secondly, a number of cell cycle genes and proteins were found to be dysregulated in FXS and *FMR1*KO neurons in our transcriptome and proteome analysis. These include, among others, DNA replication initiators and regulators of cell cycle checkpoints. Cell cycle analysis, using neural progenitor proliferation as an assay, showed increased proliferation of FXS and *FMR1*KO neural progenitors compared with control cells. In agreement with these results, studies using murine and fly models of FXS have shown that FMRP functions in the regulation of timing and proliferative capacities of neural progenitors, with loss of FMRP leading to increased neural progenitor proliferation [43–45]. These findings suggest altered cell cycle dynamics in FXS.

A neurodevelopmental feature important for neuronal connectivity is the branching and extension of neurites that develop into axons and dendrites, which precede the formation of synaptic connections [24]. Our observation of profound deficits in neurite outgrowth and branching in *FMR1*KO and FXS neurons supports previous reports showing reduced neurite development in FMRP-deficient mouse primary neurons [43, 46] as well as hESC- and hiPSC-derived neurons [20–22]. One exception is a study examining FXS neurons derived from human fetal cortical neurospheres where no difference in neurite length or branching between FXS and control neurons was found [47]. This discrepancy may be due to the small sample size (n = 1 FXS and n = 2 control post-mortem fetal cortices) and the non-isogenic settings employed.

The mechanisms underlying abnormal neurite growth phenotype have not been investigated to date but are likely to involve dysregulation of one or more developmental pathways related to axon guidance and extension. This notion is supported by our RNA-seq and proteome analysis, where we find enrichment of dysregulated genes related to axon guidance, neurite outgrowth, and cell adhesion, such as *CNTN2, ANK3, GAP43, KIF5A, EFNB3, CNTN4, UNC5A, NTNG1, DSCAM, ROBO2, PTPRT, CNTNAP2, GPRIN3, KIF5C* and *CNTN1*. As an example, *CNTN4* encodes Contactin-4, which is an axon-associated cell adhesion molecule that functions in neuronal network formation and plasticity [48]. *DSCAM* encodes Down Syndrome Cell Adhesion molecule that plays a role in neuronal self-avoidance [49]. *ROBO2* encodes for Roundabout, Axon Guidance Receptor, Homolog 2 that promotes axon guidance and cell migration. *PTPRT*, an FMRP substrate [50], is a protein tyrosine phosphatase, receptor Type T involved in synapse formation and dendritic arborization [50]. *NTNG1*, encoding for Netrin-G1 receptor, is involved in promoting neurite outgrowth of axons and dendrites [51]. *CNTNAP2*, an autism-linked gene encoding a neurexin-related cell-adhesion molecule, has been shown to influence neurite outgrowth [52, 53]. Downregulation of these key genes and associated proteins might account for the neurite outgrowth phenotype in FMRP-deficient neurons. Interestingly, some of these genes (*DSCAM, NTNG1, UNC5A, GAP43, CNTN4, CNTNAP2, GPRIN3, KIF5C*) have been associated with other neurodevelopmental disorders [54–58].

One potential consequence of neurite growth deficits is reduced neuronal network connectivity. Indeed, the significant reduction in spontaneous firings we observed using longitudinal MEA recordings in FMRP-deficient neurons is consistent with this notion. On the other hand, patch clamp recordings of neurons derived from FXS hESCs have also shown impaired ability to fire repetitive action potentials, and had reduced inward/outward currents [24]. This is consistent with our patch clamping recordings where we find a significantly reduced proportion of FMRP-deficient neurons that fire multiple action potentials compared with control. Furthermore, a well-known role for FMRP is in the regulation of key proteins involved in synaptic function and neurotransmission [59]. This is corroborated by our transcriptome and proteome analyses where we observed dysregulation of genes involved in neuronal excitability, neurotransmitter secretion, and synaptic transmission such as potassium channels in FMRP-deficient neurons. Thus, the reduced spontaneous firings we observe in FXS and *FMR1*KO neurons are likely the consequence of both impaired neuronal connectivity as well as altered intrinsic firing properties.

Our transcriptome and proteome analyses delineate dysregulated pathways affected in the absence of FMRP, highlighting mechanisms of potential relevance to the clinical manifestations of FXS. Overall, our study describes an isogenic hPSC-based model of FXS that can serve as a platform to investigate the pathophysiology of disease, and to screen and validate targets of therapeutic potential, in the context of human neurons.

## Materials and Methods

### Cell culture

HEK293 cells for assessing CRISPR/Cas9 activity were cultured in Dulbecco’s Modified Eagle’s Medium (DMEM), supplemented with 10% Fetal Bovine Serum (FBS). Control H1 hESCs (WiCell, Wisconsin) and WCMC-37 FXS hESCs (obtained from Nikica Zaninovic, Weill Cornell Medical College of Cornell University, New York) were grown using feeder-free conditions and passaged on Matrigel (BD Biosciences, Franklin Lakes, NJ)-coated plates in mTESR1 medium (STEMCELL Technologies, Vancouver, Canada). Cells were passaged by 7-min incubation with Dispase (STEMCELL Technologies, Vancouver, Canada). All hESCs have tested negative for mycoplasma.

### Generation of isogenic *FMR1* knockout (KO) lines

To create *FMR1* KO lines, CRISPR/ Cas9 was used to target the third exon of *FMR1* using a gRNA with the following sequence: 5-TTTACAGCTGGCAGCCTGATAGG-3. gRNA was cloned into pSpCas9(BB)-2A-Puro (PX459) V2.0 (Addgene plasmid # 62988), which contains a puromycin cassette to facilitate selection of transfected cells. The cloned vector was electroporated into H1 hES cells using the Neon Transfection System (Thermo Fisher) at 1,400 V for 3 pulses of 10 ms. Puromycin (1 µg/ml) was added to the culture 48 h after electroporation for a period of 48 h to enrich for transfected cells. Colonies derived from puromycin-resistant cells were picked and expanded for screening. PCR amplification using the following primers 5-TGCTTGGGAATTAGAGGGCA-3 (forward) and 5-TTGCGGCAGTGACTTTCAAA-3 (reverse), followed by the Surveyor Assay (following the manufacturer’s instructions), were used to identify correctly targeted clones. Two clones, harboring frameshift indels in *FMR1*, were selected for further characterization.

### Karyotyping

Karyotypes were determined from G-banding analysis using standard protocol according to the ISCN nomenclature. Karyotyping was performed as a service by the DNA Diagnostic and Research Laboratory at the KK Women and Children’s Hospital, Singapore.

### Testing for genomic Cas9 integration

To assay for possible genomic integration of Cas9, 100ng of purified DNA extracted from H1, FXS, KO1, KO2 along with Cas9 donor plasmid were used as template DNA and amplified with F 5’-GAAGAAGAATGGCCTGTTCG-3’ and R 5’-GCCTTATCCAGTTCGCTCAG-3’ with KOD Xtreme (Novagen, #71975). Cycling conditions were as follows: initial denaturation at 96°C for 5 mins, followed by 30 cycles of 96°C for 45s, 70°C ramped to 58°C (−0.2°C/s, total duration of 1min) and 72°C for 3mins and a final elongation at 72°C for 10mins. Amplicons were visualized on 1% agarose gel on the Geldoc XR system (Bio-Rad).

### Differentiation of three germ layers

To form embryoid bodies (EBs), hESCs were dissociated into clumps using Dispase and cultured on low-attachment tissue culture plates in KSR medium (DMEM/F12 with 20% Knock-Out Serum, 1% GlutaMax, % non-essential amino acids, and 0.1 mM 2-mercaptoethanol). Medium was changed every two days for a total of seven days. EBs were then harvested onto Matrigel-coated coverslips and left to spontaneously differentiate for nine days in KSR medium before fixation with 4% paraformaldehyde and staining.

### Neural differentiation

hESCs were induced into neural progenitor cells (NPCs) according to a previously published protocol [29]. Briefly, single-cell dissociated hESC at a density of 30,000 cells/cm2 were plated in neural induction media (NIM, DMEM/F12:NeuroBasal media 1:1 with 1% N2, 2% B27, 1% PenStrep, 1% GlutaMax, 10 ng/ml hLIF, and 5 µg/ml Bovine Serum Albumin) containing 4 µM CHIR99021 (Tocris), 3 µM SB431542 (Sigma), and 0.1 µM Compound E (Millipore) for the first seven days. The culture was then split at a 1:3 ratio for the next five passages using Accutase in NIM without Compound E on Matrigel-coated plates.

For neuronal differentiation, NPCs were plated at a density of 20,000 cells/cm2 on 50 µg/ml poly-L-ornithine/10 µg/ml laminin-coated plates and grown in NeuroDiff media (DMEM/F12/Neurobasal media (1:1) supplemented with 1% N2, 2% B27, 20 ng/ml GDNF (RD Systems), 20 ng/ml BDNF (RD Systems), 300 µM dibutyryl-cyclic AMP (D0260, Sigma Aldrich), and 200 nM L-Ascorbic Acid (A4403, Sigma Aldrich)) for at least three weeks. Medium was changed every 2-3 days.

### Flow cytometry analysis of neurons

Differentiated neurons were analysed by flow cytometry as described previously [30]. Briefly, at day 37 of differentiation, neurons were dissociated with Accutase for 10 minutes, and subsequently washed in DMEM/F12 to remove the Accutase. Dissociated neurons were treated with DNAse I for 10 minutes at room temperature and strained through a 70 µm cell strainer. Neurons were washed and resuspended in NeuroDiff sorting medium, consisting of NeuroDiff media with the addition of 0.5% BSA, 50 µg/ml Gentamycin and 0.5 mM EDTA. Cells were stained with fluorochrome-conjugated antibodies (anti-CD15-V450-#642922, anti-CD24-PE-Cy7 #561646, anti-CD44-PE #550989, anti-CD184-PE-#555974; BD Biosciences) for 30 minutes on ice. Cells were washed and resuspended in NeuroDiff sorting medium at a concentration of 2 million cells/ml. Fluorescence activated cell sorting was performed with a FACSAria II (BD Biosciences) with a 100 µm nozzle. Neurons were quantified based on the expression of CD184-, CD44-, CD15LOW, CD24+, and non-neuronal cells (glia) were quantified based on the expression of CD44+ and CD184+.

### Neurite outgrowth measurements

To assess neurite outgrowth, NPCs were plated at a density of 15,000 cells/cm2 in NeuroDiff media with modifications (substituting N2 with CultureOne supplement (Thermo Fisher)). The plate was imaged using an IncuCyte Zoom Imaging system (Essen Bioscience, Ann Arbor, MI) every 2 h for five days. For each genotype, quadruplicate live-capture measurements were performed in 9 image field per well. Cells were imaged under phase contrast, and analysis was performed using IncuCyte’s NeuroTrack module. The growth rate of neurites in each well was obtained by measuring the surface area covered by neurites and expressed as mm/mm2. The neurite outgrowth experiments have been repeated more than 3 times.

### Immunofluorescence staining

Cells were plated on ethanol-treated coverslips and fixed with 4% formaldehyde in phosphate buffer saline (PBS) for 15 min at room temperature. After washing with Tris-Buffered Saline (TBS), cells were incubated in blocking buffer (TBS containing 5% goat serum, 1% Bovine Serum Albumin, and 0.1% Triton-X-100 (Sigma Aldrich)) for 45 min at room temperature. Primary antibodies were incubated with fixed cells overnight at 4°C in blocking buffer without Triton-X-100. The following primary antibodies were used: anti-FMRP mouse monoclonal (MAB2160, Merck-Millipore), anti-OCT-4 rabbit polyclonal (sc-9081, Santa Cruz), anti-SSEA4 mouse monoclonal (MAB4304, Merck-Millipore), anti-AFP (ST1673, Merck-Millipore), anti-ASM-1 mouse monoclonal (CBL171, Merck-Millipore), anti-MAP2 rabbit polyclonal (AB5622, EMD Millipore), anti-TUJ1 mouse monoclonal (MAB1637, Merck-Millipore), anti-TBR1 rabbit polyclonal (AB31940, Abcam), anti-GABA rabbit polyclonal (A2052, Sigma), anti-PAX6 rabbit polyclonal (PRB-278P, BioLegend), anti-Nestin mouse monoclonal (MAB5326, Merck-Millipore), anti-ZO-1 rabbit polyclonal (617300, Merck-Millipore), and anti-Ki67 mouse monoclonal (MAB4190, Merck-Millipore). Cells were subsequently stained with secondary antibodies conjugated to Alexa Fluor 555 or 488 (Molecular Probes, Thermo Fisher) for 1 h at room temperature in the dark and incubated with 1 µg/ml 4’,6-diamidino-2-phenylindole (DAPI, Sigma Aldrich) for 10 min. Images were captured using an FV1000 Inverted Confocal System.

### Immunocytochemistry quantification analysis

To determine the proportion of positive cells, images were captured from at least 10 randomly selected areas. Quantification for each sample/genotype was performed blindly in a minimum of 3 coverslips. ImageJ software was then used to compute the total number of cells (DAPI-stained nuclei) and the number of cells expressing the markers. All data were expressed as the mean and standard error of the mean from three independent replicates. Statistical analysis was carried out using one-way ANOVA with Tukey’s post-hoc analysis, and significance was defined with p-value<0.05.

### Immunoblotting

Cells were lysed with RIPA buffer (Sigma Aldrich) containing cOmplete Protease Inhibitor cocktail tablets (Roche). Protein concentration was measured using the Bradford assay (BioRad). The samples were denatured at 70°C for 10 min in 4× NuPAGE sample buffer and 10× NuPAGE reducing agent (Thermo Fisher). A total of 30 µg of protein per sample was separated on 4-12% Bis-Tris gradient gels in MOPS SDS running buffer (Thermo Fisher) at 100 V for 3 h followed by transfer to nitrocellulose membrane at 120 V for 1.5 h at room temperature. The following primary antibodies were used for detection: anti-FMRP (MAB2160, Millipore; 6B8, Biolegend), and anti-Calnexin (Sigma, C4731). Alexa-Fluor 680 goat anti-mouse (Thermo Fisher) and DyLight 800 goat anti-rabbit (Rockland) were used as secondary antibodies. Membranes were imaged using the Li-Cor Odyssey infrared imaging system.

### MEA Recordings

Neurons on day 37 of differentiation were dissociated and re-plated on 0.1% polyethylenimine (Sigma Aldrich)-coated MEA plates (Axion Biosystems) in Brain-Phys media (STEMCELL Technologies, Vancouver, Canada) supplemented with BDNF, GDNF, cAMP, and L-Ascorbic Acid as previously described [37, 60]. Spontaneous activity was recorded at 37°C for 5 min every 2–3 days for 30 days using the Maestro MEA System (Axion Biosystem). Neural signals were sampled at 12.5 kHz, high-pass filtered (200 Hz – 3 kHz), and a threshold based on six standard deviation above noise levels was set (Axion Integrated Studio software (AxIS). Detected timestamps were analyzed using custom-written Matlab scripts (R2015b) as described previously [60]. Spike frequency is calculated as [spike count / experiment duration]. Mean spike frequency function removes outliers, then calculates mean ± SEM excluding channels where no activity was detected (spike number <=1). N represents the total number of channels where activity was detected from all wells of that particular condition.

### Patch clamp recording

Electrophysiological recordings were conducted on neurons cultured on glass coverslips. The experiments were performed in the blinded sample groups. The coverslips were transferred to a recording chamber in standard recording medium containing the following (in mM): 124 NaCl, 24 NaHCO3, 13 Glucose, 5 HEPES, 2.5 KCl, 2 MgSO4, 2 CaCl2 and 1.2 NaH2PO4 (310 mOsm, pH 7.4). Cells were patch-clamped with pipettes containing the following (in mM): 130 K-Gluconate, 11 KCl, 10 HEPES, 5 NaCl, 5 Na-phosphocreatine, 2 Mg-ATP, 1 MgCl2, 0.3 Na-GTP and 0.1 EGTA (pH 7.4, 300 mOsm). Action potentials are evoked by injecting depolarizing current pulses in current-clamp mode. Signals were amplified with EPC 10 USB (HEKA Elektronik, Germany) and recorded with the accompanying PATCHMASTER software. Data were analyzed using AxoGraph Ver 1.7.0 (AxoGraph Company, CA, USA) and GraphPad Prism Ver 6.00 (GraphPad Software, CA, USA).

### RNA isolation, cDNA synthesis, and Quantitative PCR

Cells were lysed using RLT Plus buffer and RNA was purified using the RNeasy Plus Mini kit (Qiagen) according to the manufacturer’s instructions. For all samples, cDNA was generated in 20 µl reactions using High-Capacity Reverse Transcriptase kit (ABI, Thermo Fisher). Quantitative real time PCR (qRT-PCR) reactions were performed in the StepOnePlus or Quant Studio6 Flex Real Time PCR System (ABI, Thermo Fisher) with ten-fold dilution of cDNA and 200 nM of each primer using the SYBR Select PCR Master Mix (ABI, Thermo Fisher). Primers are listed in **Table S1**. Relative gene expression levels were calculated using the comparative Ct method, and normalized against a control with human GAPDH.

### RNA-seq

RNA was extracted from cells on day 37 of neuronal differentiation using the RNeasy Plus Mini kit (Qiagen) according to the manufacturer’s instructions. Subsequent library preparation and paired-end 150-bp sequencing and 20 million reads/per sample using HiSeq4000 were performed by Novogene (Hong Kong). We performed RNA-seq quantification directly from the reads with Salmon v0.9.1 [61] using GRCh38 (hg38; Ensembl release 87) human reference genome. The data was imported into R using tximport [62] and analyzed for differentially expressed genes with DESeq2 [63]. Significant hits were identified using FDR < 0.05 cut-off and divided into upregulated (log2FC > 0) and downregulated (log2FC < 0) genes. The results were visualized using volcano plots. Statistical analyses for the RNA-seq and proteome data, were performed using R statistical software [64] and Python 3.6 (www.python.org). Functional enrichment analysis of significant hits was performed in clueGO version 2.5.2 [65]. For exploratory analysis, we initially normalized the read counts using the variance stabilizing transformation (VST) algorithm [66] for negative binomial data with a dispersion-mean trend. Sample-to-sample distances were calculated using the dist function [67–69] and plotted as heatmap to assess for the quality of the samples. Principal component analysis (PCA) performed in R showed a good separation of the groups, using only 2 principal components. Genes found significantly regulated in both FXS and*FMR1*KO cells, with the same direction of fold change, were considered of interest and their normalized expression visualized in the form of a heatmap.

### Preparations of samples for mass spectrometry analysis

For proteomic mass spectrometry (MS) analysis, cells were harvested on day 37 of neuronal differentiation. Sample preparation was performed as described previously [70, 71] with optimization for pellet cells. 100ul of SDC reduction and alkylation buffer were added and the mixture was boiled for 10 min to denature proteins. After cooling down, the proteolytic enzymes LysC and trypsin were added in a 1:100 (w/w) ratio. Digestion was performed overnight at 37°C. Peptides were acidified using tri-fluoro-acetic acid (TFA) to a final concentration of 0.1% for SDB-RPS binding and 20ug was loaded on two StageTip plugs. The StageTips were washed twice with 1% TFA and once with 0.2% TFA and centrifuged at 500xg. After washing the purified peptides were eluted by 60ul of elution buffer (80% acetonitrile, 19% ddH2O, 1%ammonia). The collected material was completely dried using a SpeedVac centrifuge at 45C (Eppendorf, Concentrator plus). Peptides were suspended in buffer A* (5% acetonitrile, 0.1% TFA) and afterwards mixed for 10 minutes at 1000rpm. Peptide concentrations were determined by Nanodrop (Thermo Fisher Scientific) measurement at A280nm and sample concentrations were adjusted to 1.0ug per injection.

### LC-MS/MS Analysis

All peptide samples were analyzed with nanoflow Easy-nLC 1200 (Thermo Fisher Scientific, Denmark) coupled to Q Exactive HF-X mass spectrometers (Thermo Fisher Scientific, Denmark). Peptides were separated on in-house packed column (75 m inner diameter × 50cm length) with 1.9 m C18 beads (Dr. Maisch, Germany). Column temperature was kept at 60°C. Peptide separation was achieved by 100 min gradients. Peptides were loaded with 0.1% formic acid and eluted with a nonlinear gradient of increasing buffer B (0.1% formic acid and 80% acetonitrile) and decreasing buffer A (0.1% formic acid) at a flow rate of 350 nl/min. Buffer B was increased slowly from 2% to 220% over 55 minutes and ramped to 40% over 40 minutes and then to 98%, where it was held for 5 minutes before being drop down to 2% again for column re-equilibration. Q Exactive HF-X mass spectrometer was operated in positive polarity mode with capillary temperature of 275 °C. Full MS survey scan resolution was set to 60,000 with an automatic gain control target value (AGC) of 3 ×10^6^ using a scan range of 350 1650 m/z and maximum injection times (IT) of 15ms. This was followed by a data-dependent higher-energy collisional dissociation (HCD) based fragmentation (normalized collision energy= 28) of up to 15 most abundant precursor ions. The MS/MS scans were obtained at 15,000 resolution with AGC target of 5×10^4^ and maximum injection time of 25ms. Repeated sequencing of peptides was reduced by dynamically excluding previously targeted peptides for 30 seconds.

### MaxQuant Data Processing

All data files were analyzed using the MaxQuant software suite 1.5.5.1 (www.maxquant.org) with the Andromeda search engine [72]. MS/MS spectra were searched against an in silico tryptic digest of Homo Sapiens proteins from the UniProt sequence database. All MS/MS spectra were searched with the following MaxQuant parameters for peptide identification: acetyl and methionine oxidation were searched as variable modifications and cysteine carbamidomethylation was set as fixed modification with maximal 2 missed cleavages. Precursors were initially matched to 4.5 ppm tolerance and 20 ppm. The false discovery rate (FDR) for protein and peptide matches was set to 1% based on Andromeda score, peptide length, and individual peptide mass errors. Peptide length was minimum 7 amino acids, minimum Andromeda score was 40 and maximal peptide mass was 4600Da. The second peptide feature was enabled. The match between runs option was also enabled with a match time window of 0.7 min and an alignment time window of 20 min. Relative label-free quantification (LFQ) was done using the MaxLFQ algorithm integrated into MaxQuant [73]. Protein quantification needed minimum two unique or razor peptides per protein group and minimum ratio count was set to 2.

### Proteome

Protein abundance quantitative data generated in MaxQuant was used as input for the downstream statistical analysis. The number of peptides and proteins identified in each experiment was extracted from the data and used to access the quality of the runs. During the preprocessing of the data, protein LFQ intensities were normalized by log2 transformation. Furthermore, potential contaminants and proteins identified in the decoy reverse database or only by site modification were excluded, resulting in a total of 5007 protein groups. In order to reduce the noise in the data, we filtered out proteins that did not fulfill the requirement of three valid values in at least one group. The missing values were then imputed with values randomly drawn from a downshifted Gaussian distribution of each sample’s valid values (downshift = 1.8 standard deviations, width = 0.3 standard deviations) [74]. To examine the correlation between experiments, we calculated Pearson’s correlation of the protein groups intensities and generated a heatmap. PCA was performed and visualized with 2 principal components, showing clear separation between the three groups. Differentially expressed proteins in FXS and *FMR1*KO cells were identified by Student’s t-test with Benjamini-Hochberg FDR correction for multiple hypotheses. The results were visualized with volcano plots and a cut-off of FDR < 0.05 was used to select significant hits, further divided into significant upregulated (log2FC > 0) and significant downregulated (log2FC < 0) hits. Hierarchical clustering of significant hits was carried out on the z-score protein intensities using Euclidean distance.

### Enrichment analysis and annotation networks

Functional enrichment analysis of significant hits was performed in was performed in Cytoscape 3.6 [75] clueGO app (version 2.5.2) [65], applying Fischer’s exact test and Benjamini-Hochberg FDR correction and using GOBP, GOCC, GOMF, and Reactome annotations. All genes identified in RNA-seq experiments and proteins selected for missing value imputation in Proteomics experiments, were used as background. A cut-off of FDR < 0.05 was used to identify all significantly enriched terms.

To facilitate the visualization of enriched terms in a network, we applied hierarchical level cut-offs to the enrichment search, as well as a minimum number of genes of interest per term. To reduce the redundancy of GO terms, the fusion option was also selected. GO terms, represented as nodes with sizes reflecting their statistical significance, are grouped and linked based on the similarity of their associated genes (kappa score 0.35), with the most significant term per group shown in bold and larger font.

A separate enrichment analysis for significant genes and proteins was performed against the SFARI dataset (downloaded from gene.sfari.org on 07/09/2018) [32] and FMRP targets dataset [4], using Chi2 test. An additional enrichment analysis for the genes found significantly regulated in both RNA-seq and Proteomics was also performed. In this case, common overall genes between the two data types were used as background.

### StringDB networks

Significantly regulated genes in both RNA-seq and Proteomics experiments were analyzed, using STRING database [76], to find meaningful protein-protein relationships. The resulting networks were plotted using the cluster edge betweenness function [77] to identify highly connected nodes, while the nodes were colored according to their average fold change.

### Neural rosette formation assay

hESCs were dissociated into single cells using Accutase and 4.5×10^6^ cells were seeded into AggreWell800 plates (STEMCELL Technologies, Vancouver, Canada) to form neural aggregates in STEMdiff Neural Induction Medium (STEMCELL Technologies, Vancouver, Canada). On day 5, neural aggregates were harvested and transferred into poly-L-ornithine/laminin-coated plates. On day 10, cells were fixed using 4% paraformaldehyde and stained with antibodies against ZO-1 (rabbit polyclonal, #617300, Merck-Millipore) and Nestin (mouse monoclonal, MAB5326, Merck-Millipore).

### Cell proliferation assay

Approximately 70% confluent neural progenitor cells were treated with 50 µM BrdU for 6 h, followed by fixation with 4% formaldehyde for 15 min at room temperature. For antigen retrieval, coverslips were incubated serially three times in ice-cold 1 N HCl for 10 min, 2 N HCl for 10 min at room temperature, 2 N HCl for 20 min at 37°C, and lastly in 1 M borate buffer for 10 min. Immunofluorescence staining and imaging was performed as described in the Immunofluorescence staining section using anti-BrdU (sc56258, Santa Cruz) and anti-Ki67 (MAB4190, Millipore) antibodies.

### Statistical analysis for biological assays

Samples were processed blind for assays, e.g. immunofluorescence quantitative measurements and electrophysiology. The Shapiro-Wilk test was used to test for normality. Details of the statistical tests used for the different analyses is reported in the respective figure legends. Statistical analysis was carried out GraphPad Prism v7 (La Jolla, CA, USA).

## Acknowledgments

We thank members of the Pouladi lab for helpful discussions and comments, members of the NNF-CPR Mass Spectrometry Platform for instrument support and technical assistance, and Dr. Nikica Zaninovic (Weill Cornell Medical College) for the WCMC-37 FXS hESC line. The work was partly funded by a Strategic Positioning Fund for Genetic Orphan Diseases (SPF2012/005) and SUREKids (IAF311019) from the Agency for Science Technology and Research (Singapore) to M.A.P. and a FRAXA Fellowship to K.H.U, and in part supported by a Joint Council Office grant (BMSI/15-800003-SBIC-00E) and the Novo Nordisk Foundation Center for Protein Research.

