## Supplemental Information for "Integrative analysis identifies key molecular signatures underlying neurodevelopmental deficits in fragile X syndrome"

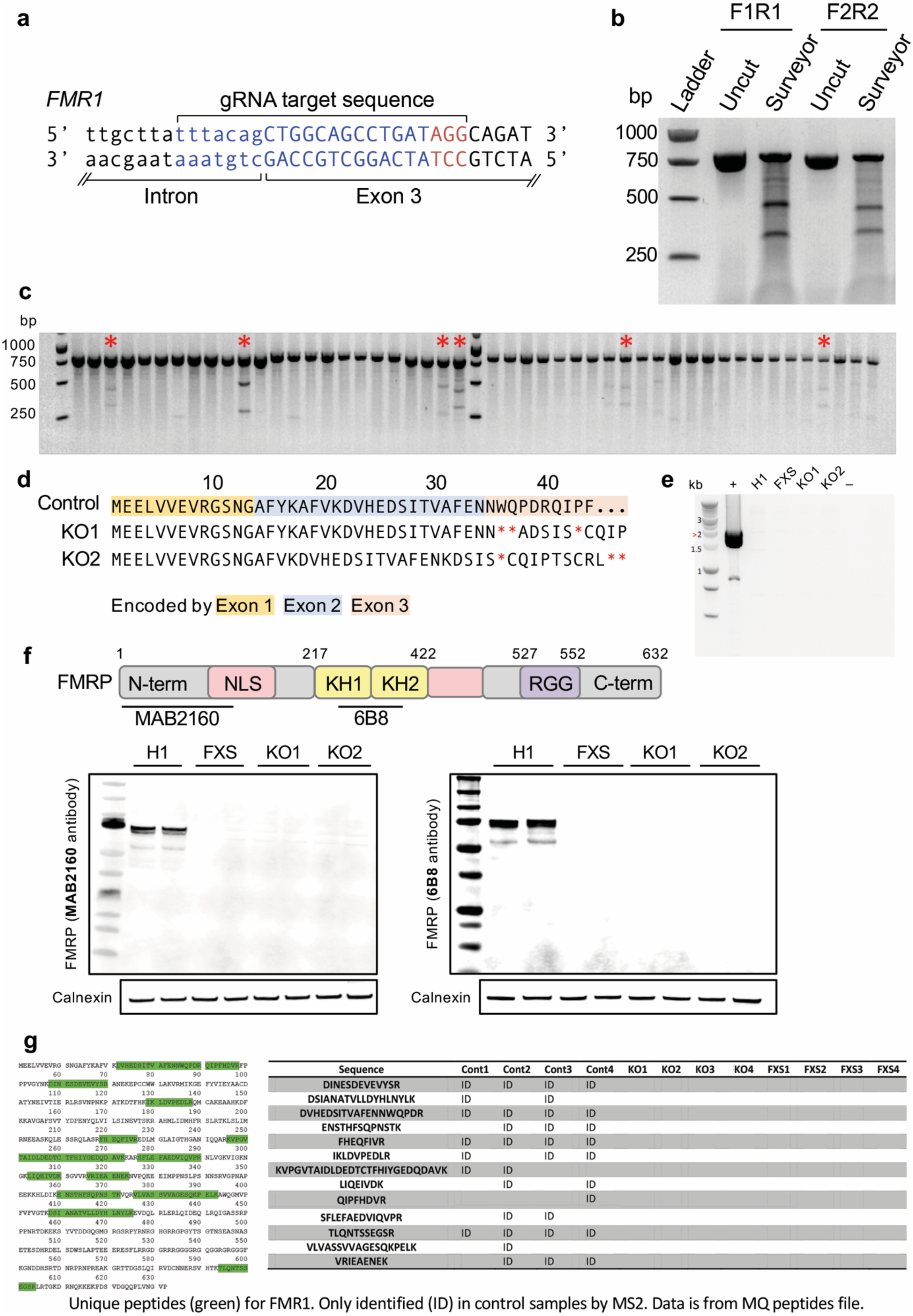


| Sequence | Cont1 | Cont2 | Cont3 | Cont4 | KO1 | KO2 | KO3 | KO4 | FXS1 | FXS2 | FXS3 | FXS4 |
|---|---|---|---|---|---|---|---|---|---|---|---|---|
| DINESDEVEVYSR | ID | ID | ID | ID | | | | | | | | |
| DSIANATVLLDYHLNYLK | ID | | ID | | | | | | | | | |
| DVHEDSITVAFENNWQPDR | ID | ID | ID | ID | | | | | | | | |
| ENSTHFSQPNSTK | | ID | ID | ID | | | | | | | | |
| FHEQFIVR | ID | ID | ID | ID | | | | | | | | |
| IKLDVPEDLR | ID | | ID | ID | | | | | | | | |
| KVPGVTAIDLDEDTCTFHIYGEDQDAVK | ID | ID | | | | | | | | | | |
| LIQEIVDK | | ID | | ID | | | | | | | | |
| QIPFHDVR | | | | ID | | | | | | | | |
| SFLEFAEDVIQVPR | | ID | ID | | | | | | | | | |
| TLQNTSSEGSR | ID | ID | ID | ID | | | | | | | | |
| VLVASSVVAGESQKPELK | | ID | | | | | | | | | | |
| VRIEAENEK | | ID | ID | ID | | | | | | | | |

**Figure S1. Targeting of *FMR1* exon 3 using CRISPR-Cas9 and verification of loss of FMRP.** (**a**) Cas9 nuclease gRNA binding sequences in *FMR1* exon 3. (**b**) Verification of activity of *FMR1*-targeted CRISPR-Cas9 using the Surveyor assay. (**c**) Surveyor screening for hESC colonies containing indels in exon 3 of *FMR1* using the F1R1 primers. Colonies marked by red asterisks were selected for single cell cloning and further screening. (**d**) Predicted truncated FMRP peptides in targeted clones. (**e**) Screening for Cas9 integration in targeted clones by PCR. (+) positive control; (–) negative control. (**f**) Verification of loss of FMRP by immunoblotting. (**g**) FMRP peptides detected by MS analysis. Unique peptides in green only detected in control neurons.

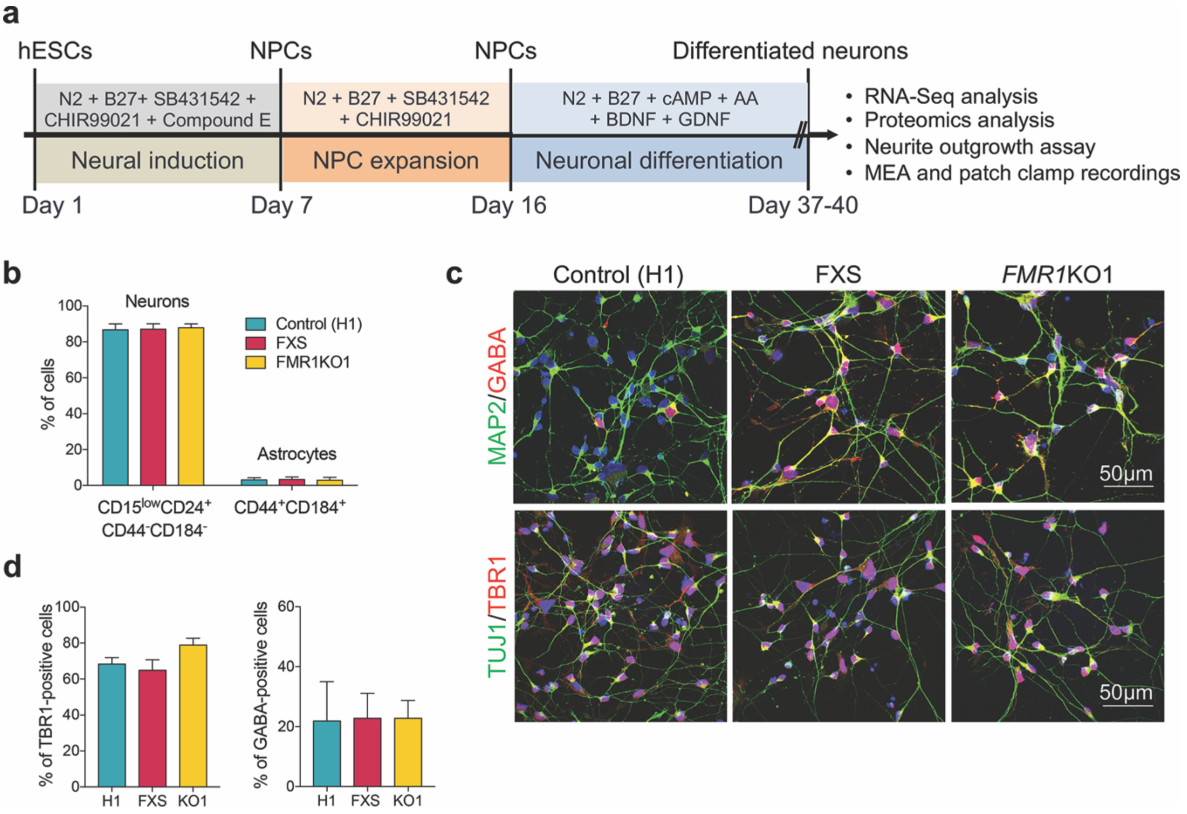


**Figure S2. Characterization of neurons derived from FXS & *FMR1*KO hESCs.** (**a**) Schematic of neuronal differentiation workflow. (**b**) Flow cytometry analysis of cellular composition following hPSC neuronal differentiation using cell surface markers of neurons and glia. (**c**) Immunofluorescence staining of MAP2/TUJ1-positive GABAergic (GABA+) and glutamatergic (TBR1+) neurons differentiated from control, FXS, and *FMR1*KO1 hESC lines. (**d**) Quantification of GABAergic and glutamatergic neurons based on anti-GABA and anti-TBR1 immunostaining.

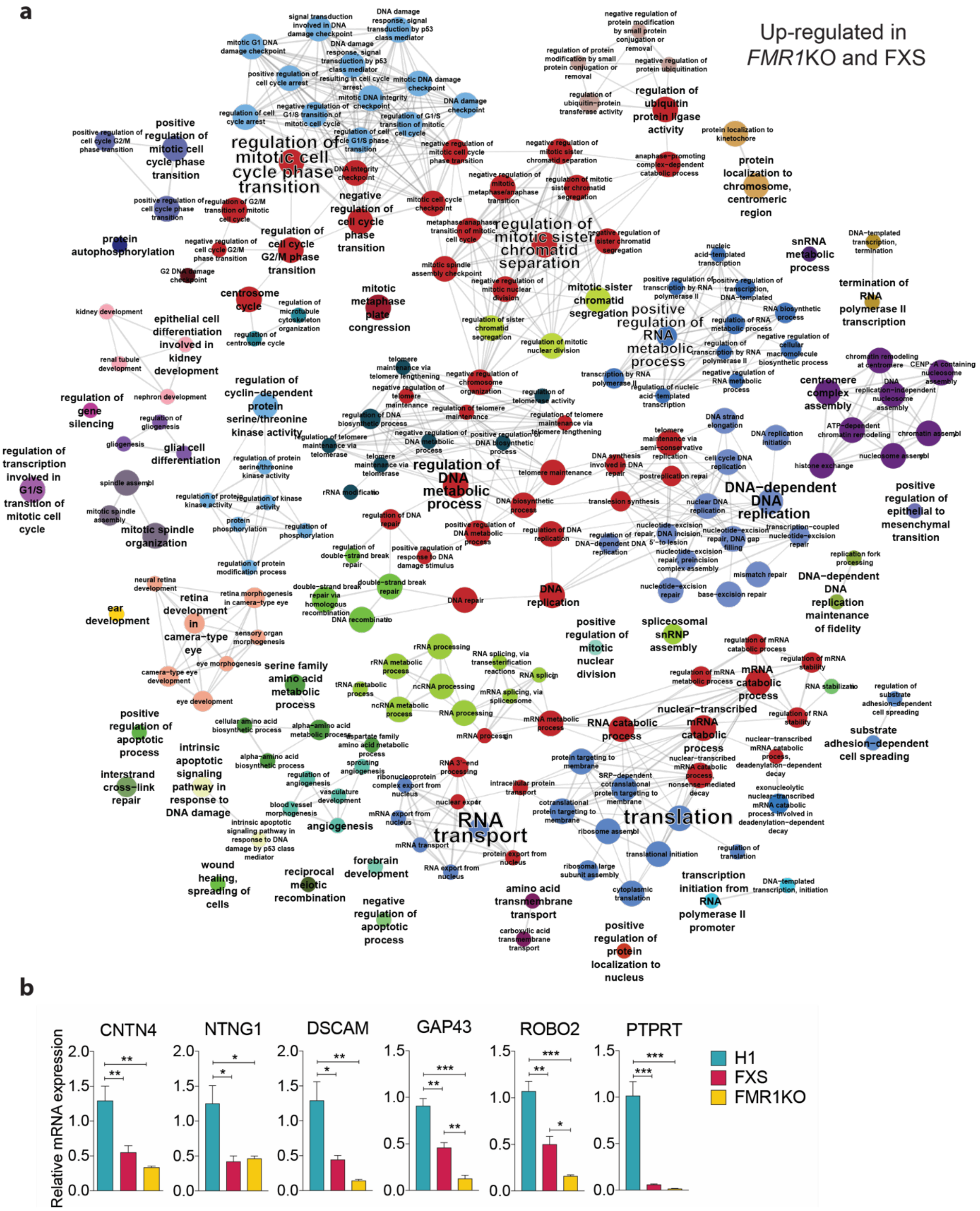


**Figure S3. Upregulated pathways and validation of select genes from the global transcriptome analysis.** (**a**) Functional annotation (biological processes) of common upregulated genes in FXS and *FMR1*KO1 neurons. **(b)** Validation by qRT-PCR of selected downregulated genes identified by RNA-seq. Values shown as mean ± SEM; n = 4 biological replicates per genotype; *p < 0.01, **p < 0.01, and ***p < 0.001 was determined by one-way ANOVA with Tukey's post-hoc test.

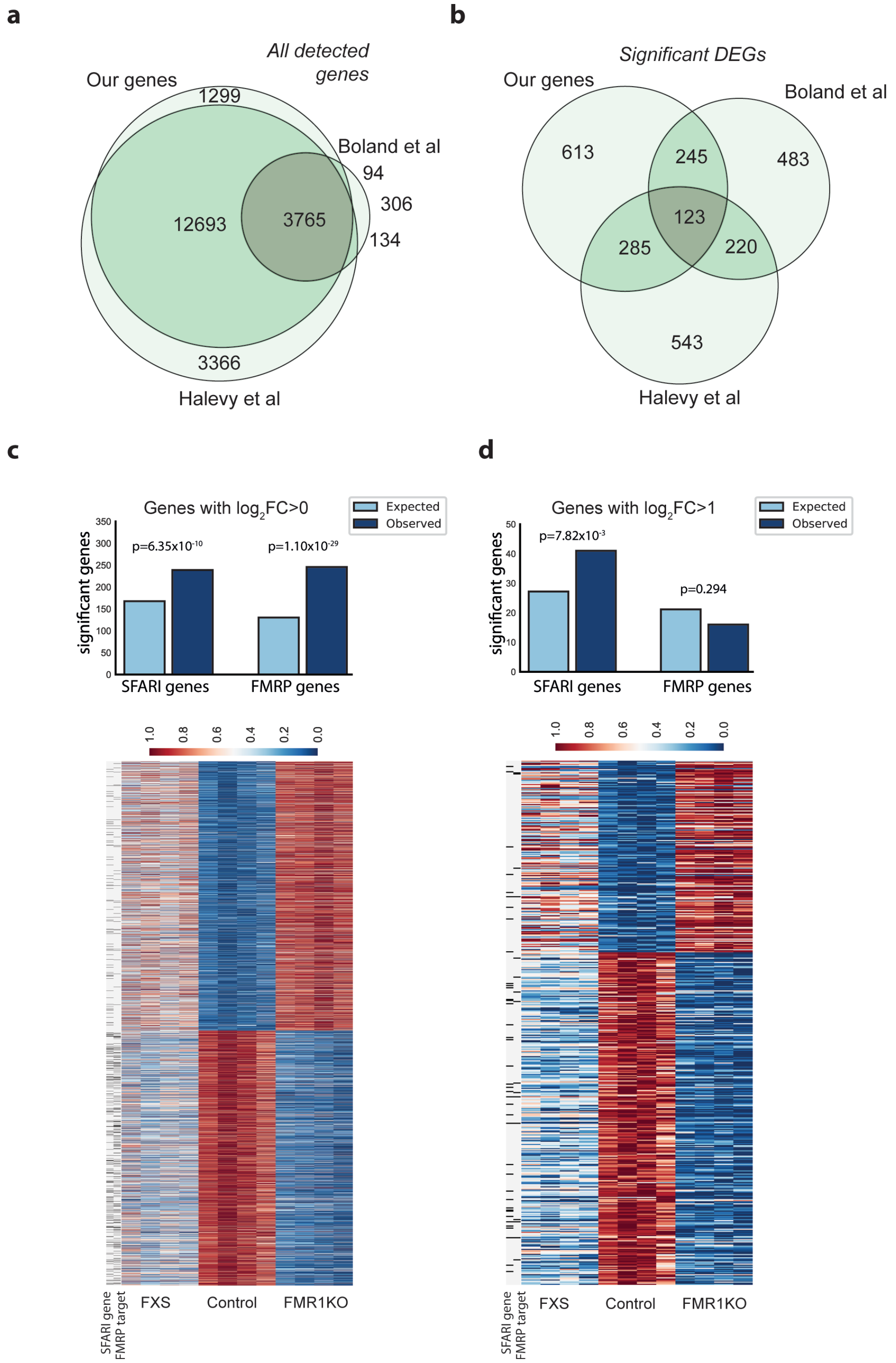


**Figure S4. Enrichment of SFARI genes and FMRP targets amongst the genes differentially expressed in FMRP-deficient neurons.** (**a**) Comparison between genes identified in this study and two others using FXS isogenic cell lines (Boland et al., 2017; Halevy et al., 2015). From the overlap (3765 genes), significantly regulated genes were compared (**b**). (**c,d**) Chi$^2$ test for enrichment of SFARI and FMRP genes amongst genes with | $\log_2$FC |>0 (**c**) and | $\log_2$FC |>1 (**d**). Heatmaps represent differentially expressed genes with SFARI and FMRP genes indicated as black lines.

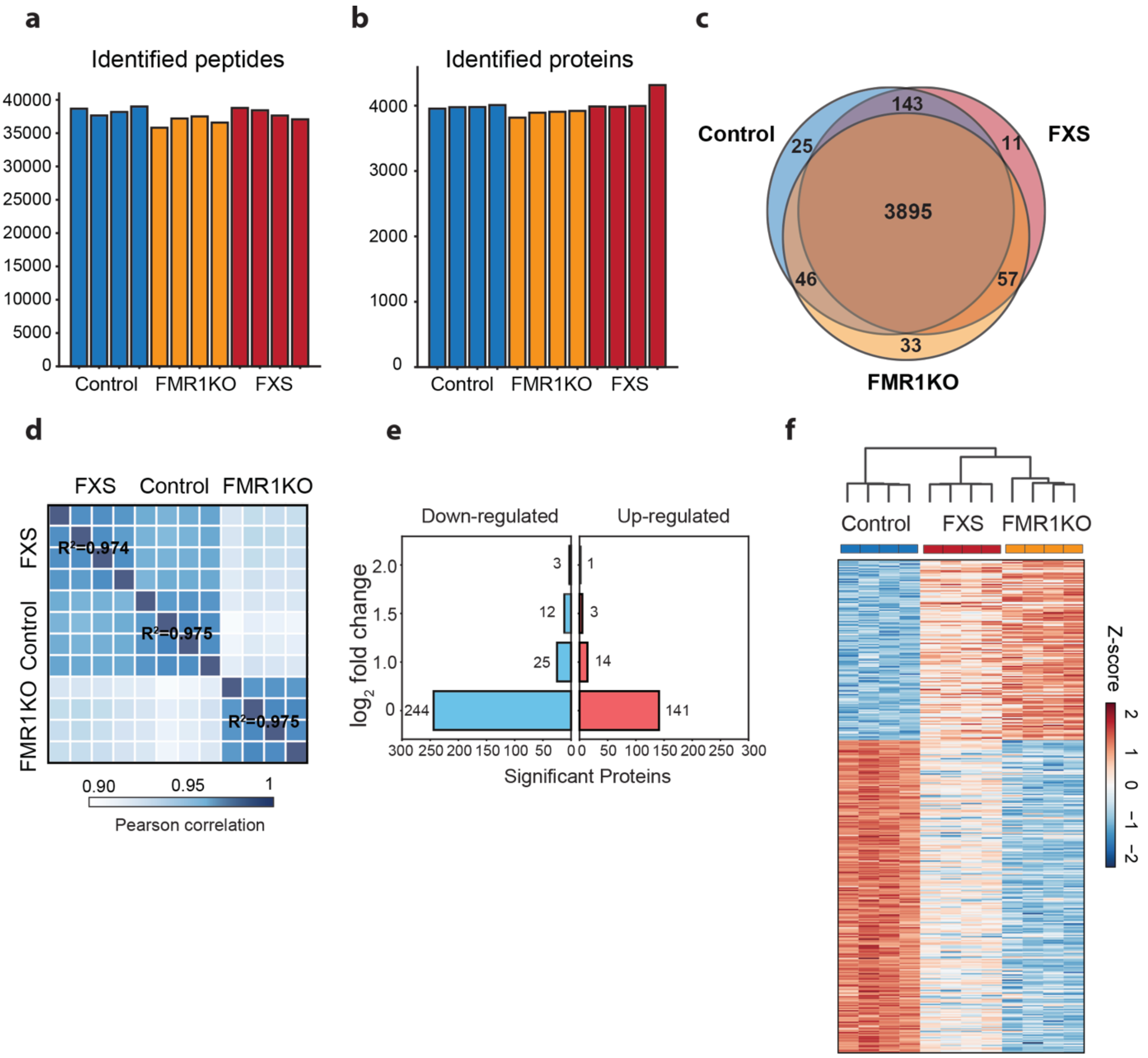


**Figure S5. Additional parameters from the global proteomics analysis.** (**a**,**b**) Number of unique peptides (**a**) and proteins (**b**) identified. (**c**) Venn diagram of proteins identified in control, FXS and *FMR1*KO neurons. (**d**) Pearson correlation of control, FXS, and *FMR1*KO replicates. (**e**) Bar plot showing the number of significant changes (down: blue, up: red) dependent on the cut-off in $log_2$ fold changes. (**f**) Heatmap and dendrogram of differentially expressed proteins.

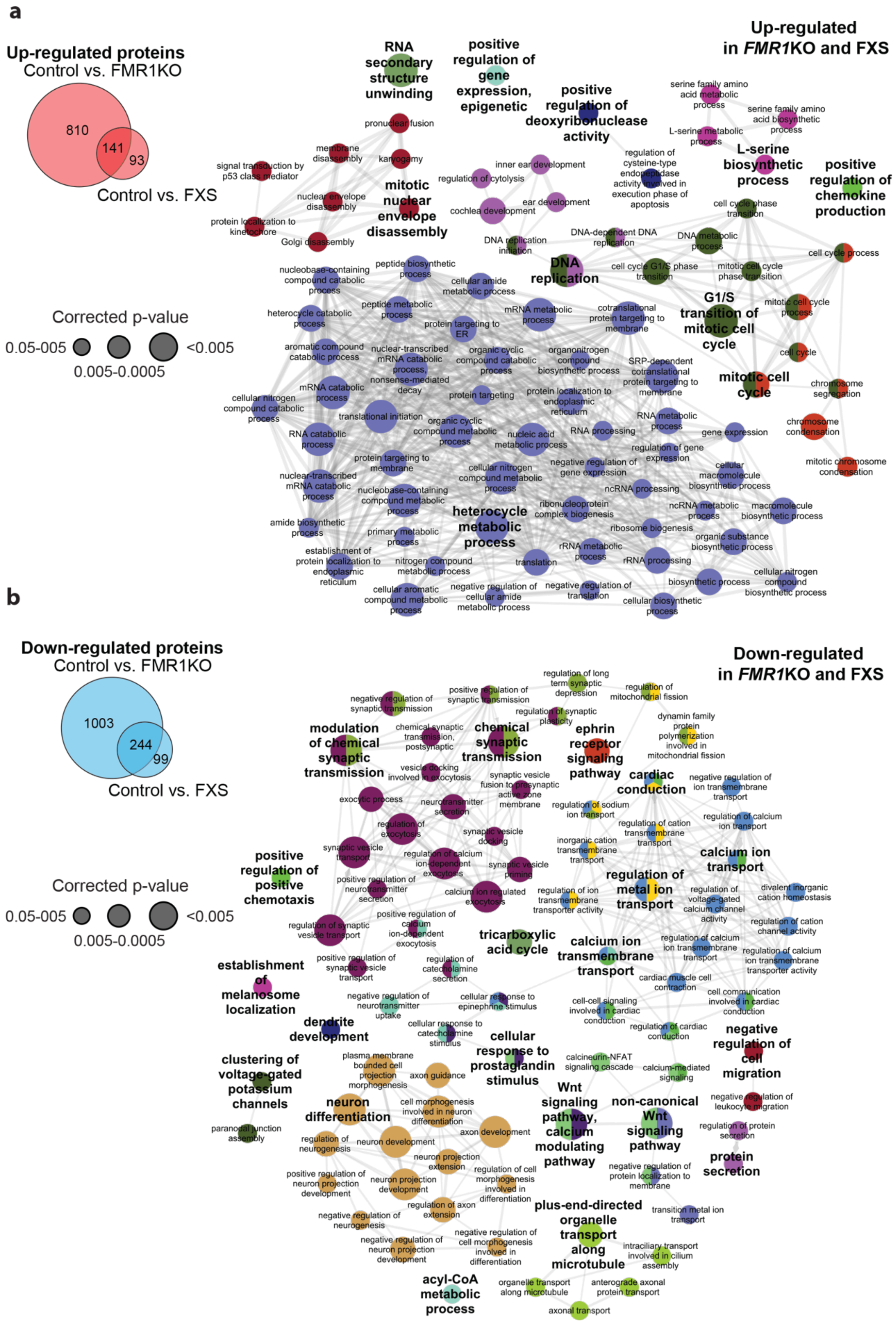


**Figure S6. Functional annotation of common protein changes in FXS and *FMR1KO* (GO biological processes), for up- (a) and down-regulated (b) proteins.** Hierarchical level cut-offs were applied to build network (**b**), a complete list of the enrichment can be found in **Table S4**.

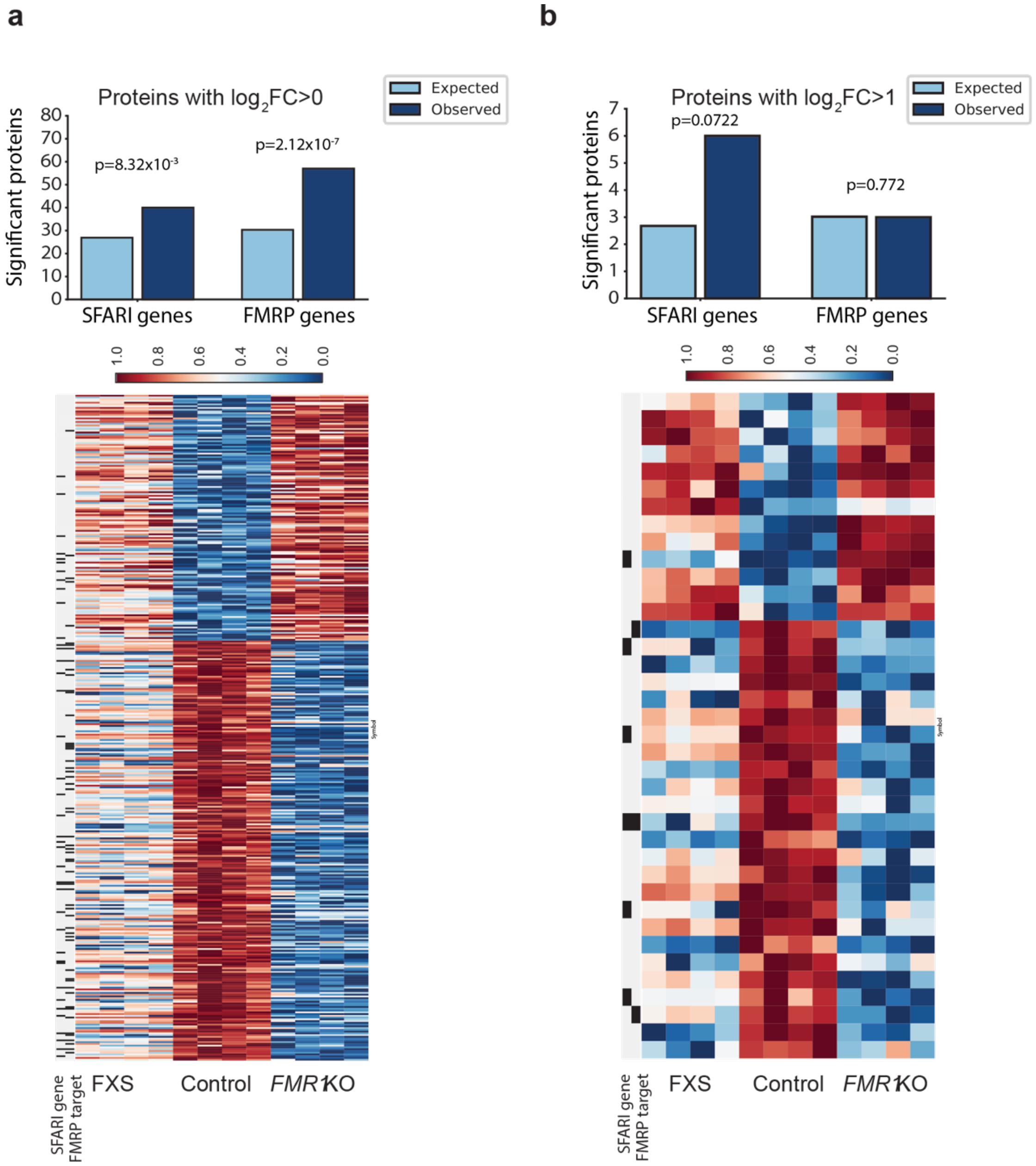


**Figure S7. Enrichment of SFARI genes and FMRP targets amongst differentially expressed proteins in FMRP-deficient neurons.** Chi$^2$ test for enrichment of SFARI and FMRP genes amongst proteins with | $log_2FC$ |>0 (**c**) and | $log_2FC$ |>1 (**d**). Heatmaps represent differentially expressed proteins with SFARI and FMRP genes indicated.

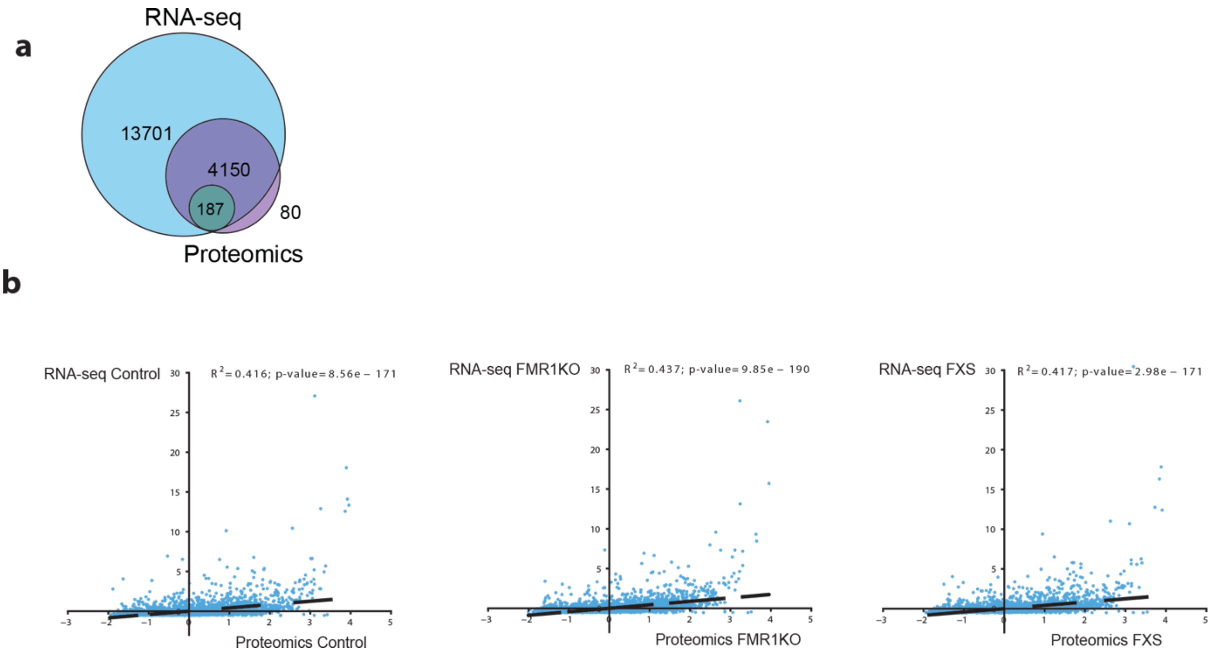


**Figure S8**. (**a**) Genes/Proteins commonly regulated in both RNA-Seq and Proteomics datasets. (**b**) Pearson's correlations between RNA-Seq and Proteomics datasets for control, FXS and *FMR1*KO datasets.

**Table S1.** Sequences of primers for qRT-PCR analysis.

| Name | F/R[a] | Sequence (5'→3') | Name | F/R[a] | Sequence (5'→3') |
|---|---|---|---|---|---|
| *OCT4* | F | AGTTTGTGCCAGGGTTTTTG | *DSCAM* | F | ACATCAAGGCTGTTTTACGGG |
| | R | ACTTCACCTTCCCTCCAACC | | R | AGATCCTGAGACAAGTGAAAC |
| *LIN28* | F | GCGGGCATCTGTAAGTGGTT | *GAP43* | F | TTCTGGATTTCAAGGGTTGAA |
| | R | GGTGAACTCCACTGCCTCAC | | R | GCTTCAGCCTCAGCAGCTTGGAC |
| *NANOG* | F | CAAAGGCAAACAACCCACTT | *PTPRT* | F | GCAGTGCCCACAGGATCTTTC |
| | R | TCTGCTGGAGGCTGAGGTAT | | R | GCTGGACCTGTCACGGCTGGAGA |
| *FMR1* | F | GTATGGTACCATTTGTTTTTGTG | *GAPDH* | F | AGCCACATCGCTCAGACACC |
| | R | CATCATCAGTCACATAGCTTTTTTC | | R | GTACTCAGCGGCCAGCATCG |

[a] Primer orientation: F, forward; R, reverse

**Table S2.** Comparison of genes found to be differentially expressed in hPSC-derived FMRP-deficient neurons (FXS and *FMR1*KO) and FXS and autism-related datasets. Significant genes (q-val<0.05 and $log_2FC>0$) were compared to SFARI genes downloaded from https://gene.sfari.org/database/human-gene/ on 07-09-2018, FMRP targets from Darnell et al., significant genes from Boland et al., and Halevy et al. ('adj.P.Val'<0.05)

| | **List of genes** |
|---|---|
| Overlap with SFARI genes | *ANK3, AP1S2, CAMK2A, CAMK2B, CELF4, CTNNB1, CUX2, DIAPH3, ARHGEF9, ARID1B, CNTN4, CNTN5, CNTN6, ASTN2, CAMTA1, ADCY5, AFF2, CREBBP, DYNC1H1, EBF3, BRCA2, BTAF1, CACNA1C, CMIP, DMD, APC, ARHGAP24, ASXL3, CASK, CHD7, ABAT, ABCA10, ABCA7, ACE, ACHE, ADARB1, ADK, CHRNA7, CNR1, ADRB2, AGAP2, CNTNAP2, APP, ASPM, CNTNAP5, ASS1, ATP2B2, ATP6V0A2, ATP8A1, ATRNL1, BAIAP2, BCL2, BRINP1, BICDL1, CACNA2D3, CADM2, CAMK4, CBLN1, CCDC88C, CDH22, CGNL1, CHD5, CHRNB3, CLSTN2, CTNND2, CUL7, DDX11, DIP2C, DNER, DOCK1, DPYSL3, DSCAM, EIF3G, ERBB4, IQSEC2, KCNQ2, MACROD2, GLRA2, KCNQ3, BIN1, CBS, FGFBP3, KHDRBS2, NRXN1, NRXN2, NRXN3, GABRA3, KCND2, KCND3, KIF5C, MAOA, MED13L, MYT1L, KMT5B, DCX, FAM19A2, FMR1, NLGN1, GABRB3, JARID2, KCNJ2, LRP2, NEXMIF, ARHGAP11B, HECW2, ITPR1, NFIA, FAT1, GABRA5, GAD1, GALNT13, GALNT14, GAP43, GAS2, GLO1, GPC4, GPR139, GPR37, GRIA1, GRID1, GRIK5, GRIP1, GRM4, ACTN4, DLGAP3, FBXO15, GRM8, H2AFZ, HECTD4, HLA-A, HLA-B, HOMER1, HS3ST5, HTR1B, IQGAP3, JAKMIP1, KIF21B, LAMA1, MAPK8IP2, MCM4, MCM6, MEGF10, MSN, MYO1E, NAV2, NCKAP5, NRG1, OPHN1, SHANK2, SLC45A1, SMARCC2, SCN1A, SLC12A5, NXPH1, PTCHD1, SATB2, SCN2A, SEMA5A, SLC4A10, STXBP1, SLC6A1, MCC, PAX6, PRICKLE1, RBFOX1, RORA, SLC6A8, SYN1, NTRK1, NTRK2, PRICKLE2, PRUNE2, ROBO2, NTNG1, NUAK1, PACS1, PAX5, PCDH8, PCDHA2, PCDHA3, PHF2, PINX1, PLCD1, PLPPR4, POLA2, PPP1R1B, PPP2R1B, PRKCB, RAB2A, RAPGEF4, RBMS3, GRIK3, GRM1, MAPK12, NELL1, PDE1C, RGS7, RIMS1, RIMS3, RPS6KA2, SAE1, SHANK1, SLC22A15, SLC30A3, SLC7A3, SLC7A5, SMC1A, SPARCL1, STK39, STX1A, SUCLG2, SYN3, TRAPPC9, TBX1, TCF20, WDFY3, XPC, TMLHE, TNIP2, TRIO, TSPAN7, PTPRT, TBC1D31, TCF7L2, WNK3, TSPAN17, UNC13A, UNC79, USP45, WNT1, ZBTB16, ZNF385B, ZNF517, TSPOAP1, RASSF5, SDK1, SERPINE1, SLIT3, SYT3* |
| Overlap with FMRP substrates | *AAK1, ABCA3, ABCG1, ACO2, ADAP1, ADARB1, ADCY1, ADCY5, ADD1, AFF3, AGAP2, AGRN, AKAP6, ALS2, ANK1, ANK3, AP2A2, APBA1, APC, APC2, APLP1, APP, ARF3, ARHGEF11, ARID1B, ARPP21, ATP13A2, ATP1A1, ATP1B1, ATP2B2, ATP2B4, ATP6V0A1, ATP6V0D1, ATP9A, ATXN1, BCAN, BMPR2, BSN, CACNB1, CADPS,* |

| | |
|---|---|
| | *CALM3, CAMK2A, CAMK2B, CAMKK2, CAMSAP1, CAMTA1, CDK5R1, CDK5R2, CELF5, CELSR2, CHD5, CHST2, CIT, CKB, CPE, CPT1C, CREBBP, CRMP1, CRTC1, CTNNB1, CTNND2, CUX2, DAB2IP, DCLK1, DDN, DENND5A, DIP2C, DIRAS2, DISP2, DLGAP3, DNAJC6, DNM1, DOCK3, DSCAM, DSCAML1, DTX1, DYNC1H1, EIF4G2, ENC1, EPB41L1, EPHA4, FAM171B, FAT1, FAT3, FBXL16, FYN, GBF1, GLUL, GNAL, GNB1, GRIK3, GRIK5, GRM4, HCN2, HERC1, HIVEP1, HIVEP2, HNRNPUL1, INPP4A, IPO5, IQSEC2, IQSEC3, ITPR1, JPH4, KALRN, KCNC3, KCND2, KCNQ2, KCNQ3, KIF1A, KIF21B, KIF3C, KIF5A, KIF5C, KLC1, LHFPL4, LINGO1, LRP8, LRRN2, MACF1, MADD, MAP1A, MAST1, MAST4, MED13L, MMP24, MTMR4, MYCBP2, MYH10, MYO18A, MYT1L, NAV1, NAV2, NAV3, NCAM1, NCDN, NCOA1, NCOA2, NCOA6, NCS1, NGEF, NRXN1, NRXN2, NRXN3, NSF, NTRK2, OLFM1, PACS1, PAK6, PCDH7, PDE2A, PHACTR1, PI4KA, PIGQ, PINK1, PITPNM1, PKP4, PLP1, PLXNA2, PPARGC1A, PPP2R1A, PPP2R2C, PPP3CA, PRICKLE2, PRKCB, PSD, PTCH1, PTK2, PTPRD, PTPRG, PTPRN2, PTPRT, QKI, RAP1GAP2, RAPGEF1, RAPGEF2, RAPGEF4, RASGRF1, RGS7BP, RHOB, RHOBTB2, RIMBP2, RTN1, RUSC2, SCN2A, SEC14L1, SEPT3, SEPT5, SGIP1, SH3BP4, SHANK1, SHANK2, SIPA1L1, SLC12A5, SLC22A17, SLC4A4, SLC4A8, SLC6A1, SLC6A17, SMARCC2, SMPD3, SNAP91, SOBP, SPARCL1, SPTAN1, SPTBN1, SPTBN2, STXBP1, SV2A, SYN1, SYNGR1, SYT1, TBC1D9, TCF20, TLN2, TMEM151A, TMEM63B, TMOD2, TNKS, TRAK2, TRIL, TRIO, TRPM3, TSHZ1, TSPAN7, TSPYL4, TTC3, TTLL7, TTYH1, TULP4, UBQLN2, UNC13A, UNC13C, UNC5A, VAMP2, WASF1, WDFY3, YWHAG, ZC3H7B, ZCCHC14, ZEB2, ZER1, ZFHX2, ZMIZ1, ZNF521, ZNF536* |
| Comparison with published datasets (Halevy et al, Stem Cell Reports 2015, and Boland et al, Brain 2017) | *ABAT, ABHD8, ABLIM3, ACAA2, ACHE, ACKR3, ACOT7, ACTA2, ACTG2, ACTL6B, ACTN1, ACTN3, ADAM15, ADAP1, ADCY1, ADCYAP1, ADD2, ADM, AES, AFAP1, AFF2, AHCY, AIF1L, AK5, AKR1C2, ALPL, ANK3, ANKRD1, ANP32B, ANXA3, ANXA5, ANXA7, AP3B2, APC2, APCDD1, APLP1, APOL2, ARHGAP24, ARHGAP36, ARHGDIB, ARHGEF16, ARHGEF39, ARL6IP5, ARPC1B, ARRDC4, ASCC3, ASCL1, ASIC1, ASIC2, ASIC4, ASPM, ASS1, ATP1B1, ATP2B4, ATP6V0C, ATP6V0D1, ATXN1, ATXN7L2, AURKA, AXIN2, B2M, B3GALNT1, BAMBI, BCAT1, BCL2L1, BCL2L12, BIN1, BIRC5, BLCAP, BLM, BMPR2, BORA, BRINP1, BSN, BST2, C12orf75, C18orf54, C1QTNF4, C1orf198, CA14, CACNA1C, CACNA2D3, CADPS, CALD1, CAMK2B, CAMTA1, CBLN1, CBLN2, CCNB1, CCNB1IP1, CCNB2, CCNC, CCNF, CD200, CD99, CDC20, CDC25A, CDC45, CDC7, CDCA5, CDCA7, CDH6, CDK1, CDK5, CDKN1A, CDKN2D, CEBPB, CELF4, CENPE, CENPF, CENPH, CENPN, CENPV, CEP152, CEP55, CERS5, CERS6, CGNL1, CHAF1B, CHD5, CHD7, CHEK1,* |

*CHEK2, CHGB, CHRNA3, CHRNA5, CHST15, CHST2, CHSY3, CISD1, CKAP2L, CKB, CKS1B, CLDN1, CLSTN2, CLTB, CMTM6, CNIH2, CNR1, CNRIP1, CNST, CNTN1, CNTN2, COL11A1, COL1A2, COL3A1, COL4A1, COL4A5, COLEC12, COMT, CPE, CPNE8, CPSF3, CPXM1, CRIM1, CRMP1, CRTAP, CRYGC, CSE1L, CSRP1, CTGF, CTIF, CTNNB1, CTNND2, CXCL12, CXCL14, CXCR4, CYB5D2, DAB2, DACT1, DCTD, DCX, DEK, DENND1A, DENND3, DEPDC1B, DERA, DHX15, DIAPH2, DIRAS2, DISP1, DLGAP5, DLK1, DNA2, DNER, DNMT3B, DOK6, DPH5, DPPA4, DPYSL3, DRAM1, DSC2, DSCC1, DSG2, DYNC1I1, DYNLRB1, E2F2, EBF1, ECEL1, ECT2, EFEMP1, EFHD1, EFNA3, EFNB2, EIF3D, EIF4G2, ELL2, ENC1, EPB41L1, EPB41L3, EPHA2, ERC1, ERC2, EVI5L, EYA2, FAM122B, FAM13B, FAM174B, FAM19A2, FAM217B, FAM89A, FANCD2, FAT1, FAT3, FAXC, FBLN2, FBXO11, FEZ2, FGF12, FGF2, FGF7, FILIP1, FLNC, FNBP1L, FNDC5, FOLR1, FOXN2, FREM2, FRMD3, FSTL1, FUCA1, FYN, FZD2, FZD5, FZD7, GABRB3, GABRE, GAD1, GALNT13, GALNT9, GAP43, GBE1, GDAP1, GFRA1, GFRA2, GINS2, GLDC, GLRA2, GNG12, GOLGA3, GPAA1, GPC4, GRIA2, GRIN3A, GSTP1, HAPLN1, HAUS4, HAUS6, HDAC1, HEBP1, HELLS, HERC1, HES5, HIGD1A, HJURP, HMGB2, HMGXB4, HMMR, HSDL2, HSPA1B, ID1, ID3, IFI6, IFITM1, IGF2BP3, INA, IRX3, IRX5, ISCA1, ISL1, ITGB1, JAG1, KCNK12, KCNQ2, KDELC2, KIF11, KIF15, KIF1A, KIF20A, KIF20B, KIF21B, KIF22, KIF26B, KIF4A, KIF5C, KLF6, KLHL1, KLHL4, KLHL42, KNSTRN, KNTC1, KRT18, KRT8, LAMP5, LAPTM4B, LGI1, LHX2, LIN28A, LIN28B, LIN7C, LINGO1, LINGO2, LIPG, LIX1, LMO3, LMX1A, LPAR4, LRAT, LRRC17, LSM14A, LY6H, LYN, LZTS1, MAB21L2, MAD2L1, MAFB, MAGEE1, MAP1A, MAP6, MAPRE3, MAPT, MASTL, MATN2, MBTPS1, MCM10, MCM2, MCM3, MCM4, MCM6, MEIS1, MEIS2, MELK, MEST, MFAP4, MFSD6, MGAT4C, MGAT5, MGME1, MGST1, MIS18A, MIS18BP1, MLLT11, MND1, MOXD1, MPC1, MRPL20, MSX1, MTMR4, MTMR7, MTSS1, MXRA7, MYO5B, MYT1L, NAP1L2, NAP1L3, NAP1L5, NASP, NAV1, NCAM1, NCAPD2, NCAPG, NDC1, NDC80, NEBL, NEDD4L, NEDD9, NEK2, NELL1, NELL2, NES, NET1, NINJ1, NKAIN4, NKD2, NMNAT2, NOTCH1, NOTCH3, NPM3, NR2F2, NRARP, NRG1, NRXN2, NSF, NSMCE4A, NTM, NTNG1, NUP107, NUP210, NUP37, NUSAP1, OIP5, OLFM3, ONECUT1, ONECUT2, OPCML, OTX2, PABPC1, PAICS, PARM1, PARP1, PARPBP, PARVA, PAX6, PBK, PCBP3, PCDH7, PCOLCE, PCP4, PCSK1, PCSK9, PDGFC, PDPN, PGM5, PHACTR3, PHGDH, PIF1, PIGZ, PITRM1, PKIA, PKIB, PKP4, PLA2G16, PLIN3, PLP1, PLS1, PLS3, PLXNB2, PNRC2, PPAT, PPP1R13B, PPP1R1B, PPP2R1A, PPP2R2C, PRC1, PRIM1, PRKCA, PRKCZ, PROM1, PRRC2A, PRSS23, PRUNE2, PSAT1, PSD2, PSRC1, PTBP1, PTPRD, PTPRG, PTPRO, PTPRR, PTPRT,*

*PTPRZ1, PURB, RAB11FIP1, RAB2A, RAB3B, RAB3C, RAD51AP1, RAI14, RAN, RAP1GAP2, RAPGEF5, RASA3, RBPMS2, RCC2, RCN1, REEP1, RGMA, RGS20, RGS7, RHBDL3, RHOC, RHOU, RIMBP2, RIMS2, RIMS3, RIPPLY3, RNF150, ROBO2, RORA, RPA1, RPL18A, RPL8, RPS2, RPS6KA2, RRAGD, RSPO3, RTKN, RTN1, RUNDC3A, RUNDC3B, RUSC2, SALL4, SARS2, SAT1, SCARB2, SCG3, SCN2A, SEMA3C, SEMA4D, SEMA5A, SEMA6A, SEPHS1, SERINC1, SERINC3, SERP2, SERPINI1, SFRP2, SGIP1, SH3BP4, SH3GL3, SHMT2, SHROOM2, SHROOM3, SHROOM4, SKA3, SLC12A4, SLC15A3, SLC16A14, SLC17A6, SLC1A3, SLC37A1, SLC37A4, SLC5A7, SLC7A3, SLC7A5, SLC9A3R1, SLIT2, SMC4, SNAI2, SNAP91, SNCA, SOBP, SORBS3, SOX2, SOX21, SOX9, SPNS2, SPOCK2, SPP1, SPTBN1, ST3GAL1, ST8SIA3, STMN2, STOML1, STX1B, STXBP1, SUCLG2, SULF1, SUMF1, SYBU, SYP, SYT1, TAF15, TAGLN, TAOK3, TAP2, TBC1D9, TCEAL2, TCERG1L, TDP1, TEK, TERF2IP, TFPI2, TGIF2, THSD7B, TICRR, TIMELESS, TJP2, TLE1, TLE4, TM2D3, TM7SF3, TMEFF2, TMEM123, TMEM179, TMEM243, TMEM63B, TMOD2, TMPO, TMSB15A, TMX4, TOP2A, TP53I11, TPBG, TPD52L1, TPM2, TRAM1, TRH, TRIB2, TRIM22, TRIM71, TRPM3, TSPAN12, TSPAN18, TSPAN7, TTC9, TTF2, TTK, TUBA1C, TUBB, TWSG1, TYMS, TYW3, UBE2C, UBE2T, UCP2, UNC13A, UNC13C, UNC5A, USP44, UST, VASH2, VASN, VCAM1, VCAN, VIM, VOPP1, VSTM2B, VWC2, WDHD1, XKR4, XPOT, YAP1, YBX3, YWHAG, ZCCHC12, ZDHHC22, ZEB2, ZFP30, ZFP36L1, ZMAT3, ZNF217, ZNF248*

---

**Table S3**. Comparison of proteins found to be differentially expressed in hPSC-derived FMRP-deficient neurons (FXS and *FMR1*KO) and FXS and autism-related datasets. Significant proteins (q-val<0.05 and $log_2FC>0$) were translated into codifying genes and compared to SFARI, FMRP targets from Darnell et al., significant genes from Boland et al., and Halevy et al. ('adj.P.Val'<0.05)

| | **List of genes** |
|---|---|
| Overlap with SFARI genes | *ANK2, ANK3, ASS1, ATP1A3, BAIAP2, BIN1, CD44, CHD7, CNTNAP2, DDX3X, DNM1L, DPYSL3, DYNC1H1, EBF3, FGFBP3, FMR1, GAP43, GRIP1, GTF2I, HECTD4, KIF5C, MAP2, MCM4, MCM6, MYO5A, NRXN2, PCCA, PCCB, PDE4B, PLXNA3, PPP2R1B, RAB2A, SAE1, SNAP25, SRGAP3, SSRP1, STX1A, STXBP1, SYN1, TOP* |
| Overlap with FMRP substrates | *ACO2, AGRN, ANK2, ANK3, AP2A2, ARHGEF12, ATP1A3, ATP2B4, ATP6V1B2, CADPS, CLASP2, CLTC, CPLX2, CRMP1, DCLK1, DNM1, DYNC1H1, EPB41L1, GNB1, GPRIN1, INPP4A, ITSN1, KALRN, KIF5A, KIF5C, LINGO1, LLGL1, MAP1A, MAP1B, MAP2, MYO5A, NCAM1, NRXN2, NSF, PDE4B, PLEC, PLXNA1, POLR2A, PPP1R9B, PPP3CA, QKI, RTN4R, SEPT3, SEPT5, SNAP25, SNAP91, SPTAN1, SPTBN1, SPTBN2, SRGAP3, STXBP1, SV2A, SYMPK, SYN1, SYT1, TTYH3, TUBB3* |
| Comparison with published datasets (Halevy et al, Stem Cell Reports 2015, and Boland et al, Brain 2017) | *ANK3, REEP1, RAB2A, CNTN1, TAGLN3, CHD7, NTM, NSF, CRMP1, NEFM, ASS1, GLDC, DLK1, GAP43, COMT, NCAM1, MCM3, MCM6, AHNAK, PHGDH, ATP2B4, MAPT, ANXA5, SRSF6, SCAMP1, MCM4, ACSL3, UHRF1, SNAP25, STX1B, FARP1, YWHAZ, RPS2, TP53I11, GPRIN1, MCM2, DDT, MAP6, EPB41L1, SMC4, PDE4B, MAP1B, DCAF7, RUFY3, CDK1, RPL14, AKR1C2, NRXN2, CNTN2, STMN2, SPTBN1, SLC17A6, TOP2A, SNCA, FLNC, DUT, MAB21L2, BIN1, SLC25A1, STXBP1, VIM, MYO5A, L1CAM, EDIL3, RAB3C, LINGO1, SYT1, PABPC1, GNG12, BCAT1, IDH2, ARHGEF12, NES, ATP6V1E1, SHROOM2, AFAP1, DEK, IGF2BP3, DPYSL3, ITSN1, MAP2, KIF5C, CADPS, SRGAP3, CD200, AHCY, CRIP2, EPB41L3, UGP2, TUBB3, DPYSL4, CSE1L, MAP1A, TCEA1, PCCB, NUP210, ISYNA1, PSAT1, TPBG, ARL6IP5, CCDC88A, SNAP91, NCAPD2* |

**Table S4.** Intrinsic properties of hPSC-derived control and FMRP-deficient neurons

| Parameter | Mean ± S.E.M. Control (n = 79) | FXS (n = 82) | FMRP1KO (n = 67) | P-value# |
|---|---|---|---|---|
| Average maximum no. of AP | 4.09 ± 0.64 | 1.94 ± 0.56 | 3.76 ± 0.95 | a: p = 0.001**<br>b: p = 0.094<br>c: p = 0.30 |
| Input resistance (GOhm) | 0.433 ± 0.02 | 0.400 ± 0.02 | 0.395 ± 0.02 | a: p = 0.11<br>b: p = 0.21<br>c: p = 0.74 |
| Resting membrane voltage (mV) | -61.3 ± 2.02 | -64.8 ± 1.75 | -66.2 ± 1.55 | a: p = 0.19<br>b: p = 0.15<br>c: p = 0.99 |
| AP amplitude (mV) | 53.1 ± 2.18 | 39.2 ± 2.37 | 48.7 ± 3.33 | a: p < 0.001***<br>b: p = 0.25<br>c: p = 0.02* |
| Rheobase (pA) | 36.7 ± 4.21 | 66.5 ± 7.99 | 43.4 ± 4.29 | a: p < 0.001***<br>b: p = 0.072<br>c: p = 0.11 |
| AHP (mV) | 15.8 ± 0.93 | 11.9 ± 0.98 | 14.3 ± 1.14 | a: p = 0.007**<br>b: p = 0.29<br>c: p = 0.14 |
| AHP latency (ms) | 14.2 ± 0.91 | 11.4 ± 0.64 | 13.9 ± 1.10 | a: p = 0.026*<br>b: p = 0.67<br>c: p = 0.11 |
| Half-width (ms) | 2.70 ± 0.16 | 2.91 ± 0.17 | 3.30 ± 0.34 | a: p = 0.19<br>b: p = 0.31<br>c : p = 0.81 |
| AP threshold (mV) | -27.9 ± 0.95 | -23.5 ± 1.19 | -28.7 ± 1.11 | a: p = 0.005**<br>b: p = 0.586<br>c: p = 0.002** |
| Latency to AP (ms) | 53.4 ± 10.50 | 20.1 ± 2.97 | 60.0 ± 16.70 | a: p = 0.002**<br>b: p = 0.66<br>c: p < 0.001** |

Values ± SEM, values and SEM are corrected to 3 significant figure and 2 decimal places respectively. Student's t-test was used for AP amplitude analysis while Mann-Whitney post hoc analysis was used for all the other variables. # a: Comparison between Control and FXS groups, b: Comparison between Control and *FMR1*KO groups, c: Comparison between FXS and *FMR1*KO groups; AP, action potential; AHP, Afterhyperpolarization.
